# General Purpose AI for Public Data Analysis and Experimental Hypothesis Generation

**DOI:** 10.64898/2026.10.16.759709

**Authors:** Gyuhyung Jin, Xiaoping Bao

## Abstract

Public molecular datasets can support biological questions beyond those addressed in the original studies. We examined how general-purpose AI could help generate and investigate such questions through four complementary investigations. Interactive single-cell analysis identified Apelin signaling as a candidate for promoting mesendoderm differentiation of human pluripotent stem cells, although subsequent experiments did not establish an effect under the tested conditions. Autonomous analyses then searched public single-cell data for tumor antigens that could be targeted by engineered immune cells. Three multiple-myeloma runs shared 50 candidates and repeatedly recovered BCMA and CD38, whereas three pancreatic ductal adenocarcinoma runs produced different exploratory results and nominated no target meeting their selection criteria. To examine whether published datasets could yield additional biological insights, we conducted two separate AI analyses of anonymized lung single-cell data without providing the original publication. One analysis was performed without information about the tissue type or disease status, while the other was provided with this information. Without tissue or disease information, the AI identified lung cell populations and inferred disease-associated tissue remodeling. The analyses also generated additional biological questions, eight of which showed statistical support in an independent patient dataset, including coordinated antioxidant-related expression across epithelial cell types and increased fibroblast CRYM expression. Finally, a broader public-data analysis integrated RNA expression, proteomics, CRISPR-based functional screening, and normal-tissue expression to prioritize 213 potential therapeutic targets for pancreatic cancer. Combining these measurements helped distinguish candidates with similar RNA evidence and assess their functional relevance and expression in normal tissues, although therapeutic selectivity remains to be established. Together, these investigations show how general-purpose AI can extend an experimental biologist’s use of public data, generate testable hypotheses and identify biological relationships for further study. They also show why completing an analysis and obtaining a reproducible expression pattern are distinct from establishing an experimental or therapeutic effect.

## 1. Introduction

Single-cell RNA sequencing (scRNA-seq) has made it possible to examine cell populations and molecular states within developing tissues, diseased organs and tumors. Once deposited in public repositories, these measurements provide opportunities to investigate additional biological relationships and generate hypotheses beyond the objectives of the original studies. The growing availability of standardized public single-cell datasets further expands the opportunities for such investigations [1]. Reanalysis involves decisions about suitable studies, computational methods, cell identities, biological comparisons and interpretation, all of which can influence the resulting conclusions.

Language-model-based tools can assist with several stages of this process. Single-cell systems such as CellAgent, CellAtria and CellVoyager support natural-language-guided workflows, data preparation or autonomous analysis, while scChat connects interactive single-cell exploration with biological interpretation and experimental planning [2–5]. CellWhisperer enables conversational exploration by learning relationships between expression profiles and text descriptions, including applications to biological questions in public datasets [6]. These systems address related needs through different combinations of analysis software, execution environments and user interaction. Broader biomedical agents, including Biomni and BioMedAgent, extend tool use across data types and research tasks, and GeneAgent uses biological databases to check gene-set interpretations [7–9]. Together, these studies provide a foundation for investigating how AI can help researchers connect biological questions to computational evidence.

Recent studies have also examined AI-generated hypotheses and their evaluation. Co-Scientist and Robin connect computational hypothesis generation with experimental validation performed by scientists, whereas the Virtual Biotech integrates biological and clinical evidence to investigate therapeutic questions [10–12]. Studies of autonomous and copilot analysis, together with bioinformatics benchmarks, show the importance of assessing both completed tasks and the scientific quality of their outputs [13–15]. Public-data reanalysis is particularly relevant because a computationally useful result may be a reproducible relationship within existing measurements, even before its underlying mechanism is experimentally tested. The related approaches and the scope of their reported evaluations are summarized in Supplementary Table S10.

Building on these applications of AI in biological research, we examined general-purpose AI interfaces within an experimental stem cell research workflow. Our interest was in the findings generated, their consistency across analyses and their value for developing and evaluating biological hypotheses. We used two approaches that differed in the level of user involvement and the role of AI in the analytical process. In the interactive approach, AI provided stepwise guidance for dataset selection, computational analysis, and interpretation. We executed the suggested code, examined the outputs, and discussed the findings with AI before proceeding to subsequent steps. In the autonomous approach, an AI agent received a research prompt defining the biological objectives and general requirements, then independently selected datasets, established its computational environment, performed analyses, and generated reports with minimal user intervention. By examining these two approaches, we sought to understand how general-purpose AI could support the progression from biological questions to computational findings, as well as how consistently autonomous analyses could identify biological candidates across separate runs.

We organized the study into four complementary investigations (Fig. 1). First, we examined whether interactive AI guidance could help identify a candidate signaling factor from public developmental single-cell data and translate the resulting hypothesis into an experiment using human pluripotent stem cells (hPSCs). Second, we evaluated whether an AI agent could autonomously perform tumor-antigen discovery, from dataset selection to candidate identification, and whether repeated analyses would produce consistent results. Third, we investigated whether AI could identify additional biological relationships in published single-cell data without access to the original study, comparing analyses performed with and without information about the tissue type and disease status. We also examined whether the resulting findings could be supported in an independent patient dataset. Finally, we explored how integrating RNA expression with protein measurements, functional screening, and other public data could influence therapeutic target prioritization. Together, these investigations allowed us to examine different applications of general-purpose AI in biological data analysis, from experimental hypothesis generation to autonomous analysis and integration of multiple data types.

**Figure 1.**
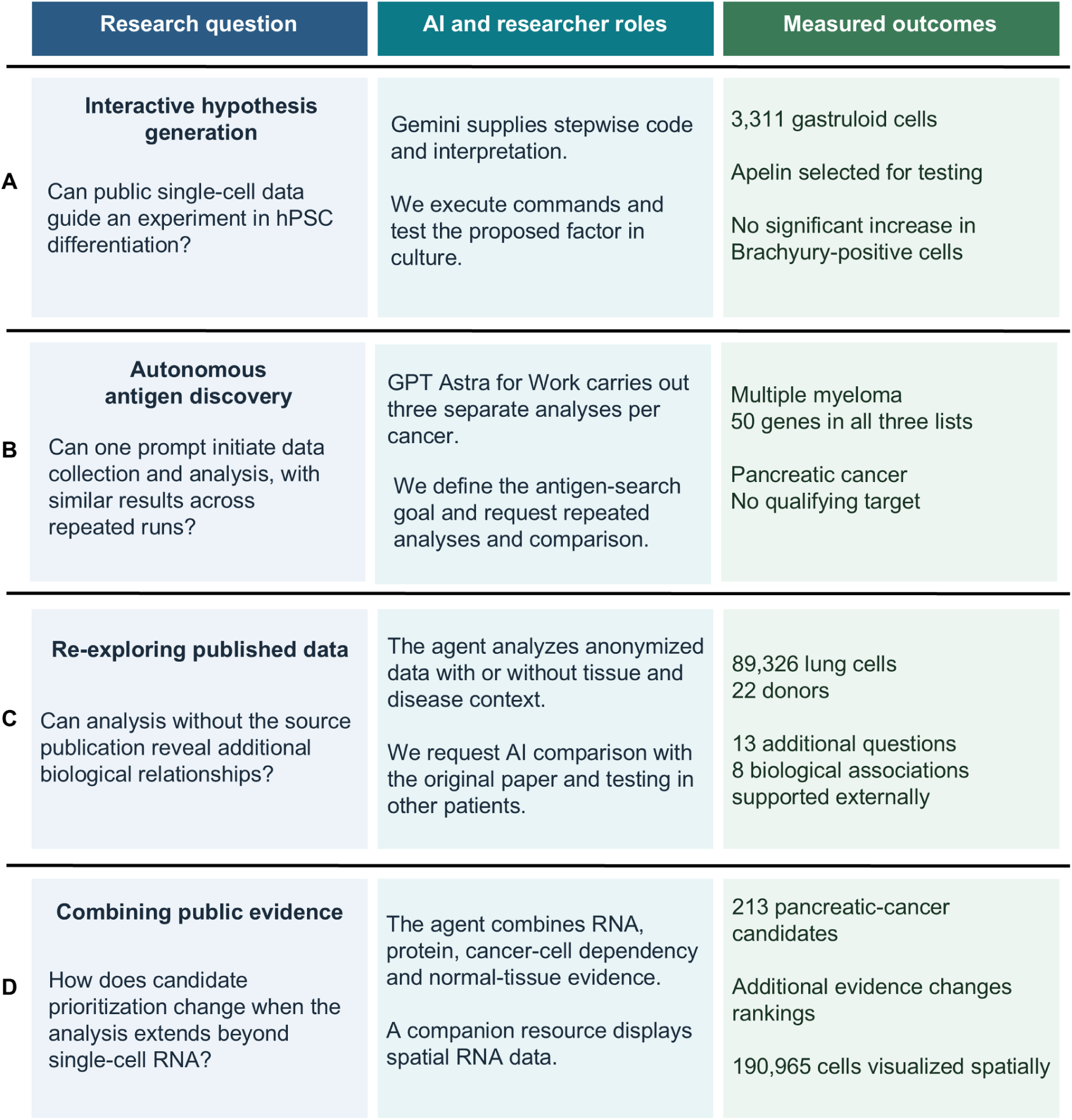
Study design and the roles of general-purpose AI in public-data analysis. (A) Interactive analysis of public gastruloid data followed by experimental testing of an Apelin hypothesis in H9 human pluripotent stem cells (hPSCs). Gemini Pro 3.1 provided stepwise guidance and code, which we executed and discussed before testing the proposed treatment. (B) Autonomous searches for tumor antigens that could be targeted by engineered immune cells. GPT-6 Astra, using the Extra high or Ultra reasoning setting, performed three separate runs for pancreatic ductal adenocarcinoma (PDAC) and multiple myeloma (MM), from data acquisition through candidate assessment. Fifty MM genes were identified in all three runs. (C) Re-exploration of anonymized lung single-cell data with or without tissue and disease information. Findings were compared with the source publication and additional biological questions were tested in other patients. (D) Integration of RNA, protein, cancer-cell dependency and normal-tissue evidence for PDAC candidate prioritization. A separate companion resource displayed spatial RNA measurements from a pancreatic tissue section. GPT-6 Astra performed the analyses in B–D. Counts summarize the main datasets and outcomes.

## 2. Results

### 2.1 Interactive single-cell analysis identified an Apelin hypothesis for hPSC mesendoderm differentiation

The first investigation used interactive AI guidance to connect public scRNA-seq data with a candidate for experimental testing. Our previous work on cardiac and immune cell differentiation from hPSCs motivated our interest in early mesodermal lineage specification [16–18]. Mesendoderm differentiation of hPSCs provided a tractable setting because extracellular signals can be added directly to the culture and early differentiation can be assessed by Brachyury expression. We asked Gemini to identify signals associated with primitive-streak and mesendoderm formation, prioritizing candidates beyond those commonly used in differentiation protocols. The instructions required stepwise guidance, inspection of the actual analysis outputs, and a distinction between gene-expression observations and proposed mechanisms. We executed the suggested commands, returned the outputs, and refined the analysis through successive exchanges.

The analysis used a published scRNA-seq dataset of micropatterned human embryonic stem cell gastruloids [19]. The samples comprised two gastruloid replicates collected after 44 h of BMP4 treatment and a separate population collected after dissociation and reseeding. These endpoint samples allowed comparison of cell states within differentiated cultures. Of 3,396 input cells, 3,311 passed quality-control filters and formed 13 clusters (Fig. 2A). The Gemini-guided analysis examined POU5F1, T (TBXT), MESP1 and SOX17 expression to identify populations relevant to pluripotency and early lineage specification (Supplementary Fig. S1). Six clusters were grouped as putative pluripotent, primitive-streak-like and mesoderm populations, containing 1,023, 562 and 496 cells, respectively (Fig. 2B).

**Figure 2.**
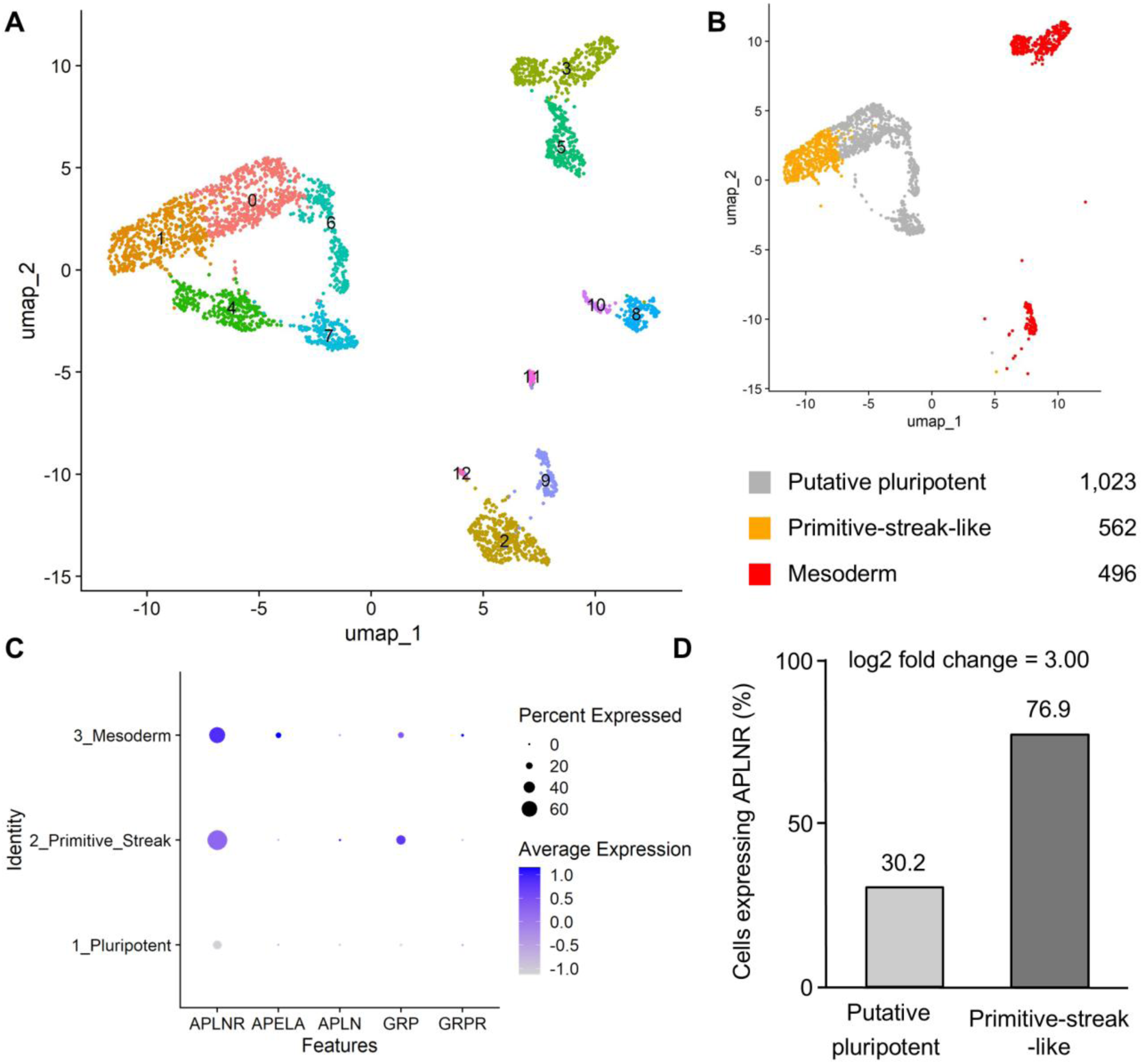
Interactive analysis of human gastruloid single-cell data identified APLNR as a candidate signaling receptor. (A) Two-dimensional map of 3,311 cells after quality control, generated using uniform manifold approximation and projection (UMAP). Each point represents one cell; nearby cells have similar gene-expression profiles, and colors indicate clusters. Samples comprised gastruloids collected after 44 h of BMP4 treatment and cells collected 48 h after dissociation and reseeding. (B) Selected cell states displayed at the same coordinates: putative pluripotent cells (1,023), primitive-streak-like cells (562) and mesoderm cells (496). (C) Candidate ligand and receptor expression in the selected cell states. Larger dots indicate a higher proportion of cells with detectable RNA, and color indicates each gene’s scaled average expression across the three states. (D) Percentage of cells with detectable APLNR, which encodes the Apelin receptor. APLNR expression was detected in 30.2% of putative pluripotent cells and 76.9% of primitive-streak-like cells; the average log2 expression difference was 3.00. Data were obtained from GSE144897 [19].

Comparison of the primitive-streak-like and putative pluripotent populations identified increased expression of developmental genes, including MESP1, MIXL1, TBX6 and PDGFRA. APLNR, which encodes the Apelin receptor, was also enriched, with an average log2 fold change of 3.00 and detectable expression in 76.9% versus 30.2% of cells (Fig. 2C,D). In contrast, the receptor ligands APLN and APELA were detected in relatively few cells. This pattern led Gemini to propose supplementing the culture with an APLNR ligand to test whether receptor activation could promote mesendoderm differentiation. The biological rationale was therefore based on receptor expression and the availability of a soluble ligand, rather than a demonstrated requirement for Apelin signaling in this system.

Prior studies provided additional context for this hypothesis. Apelin exposure has been examined during hPSC differentiation, including hematopoietic, cardiac and mesodermal differentiation [20–23]. In particular, Apelin-13 treatment during CHIR99021-containing H9 differentiation increased Brachyury mRNA at a later time point [23]. Thus, the relevant experimental question was whether Apelin could promote the early mesendoderm response in our culture conditions, including in the absence of a conventional induction signal.

We treated H9 cells with 10, 50 or 500 nM [Pyr¹]-Apelin-13 on days 0 and 1 and measured Brachyury expression by flow cytometry on day 2 (Fig. 3A). Untreated cells and cells treated with 6 µM CHIR99021 served as negative and positive controls, respectively. Apelin treatment alone resulted in low percentages of Brachyury-positive cells across all tested concentrations, whereas CHIR99021 induced Brachyury expression in approximately 86% of cells. No significant differences were observed between the untreated and Apelin-treated groups (one-way ANOVA, P=0.1952), although the 10 nM Apelin group showed a slightly higher mean with substantial variation among replicates.

**Figure 3.**
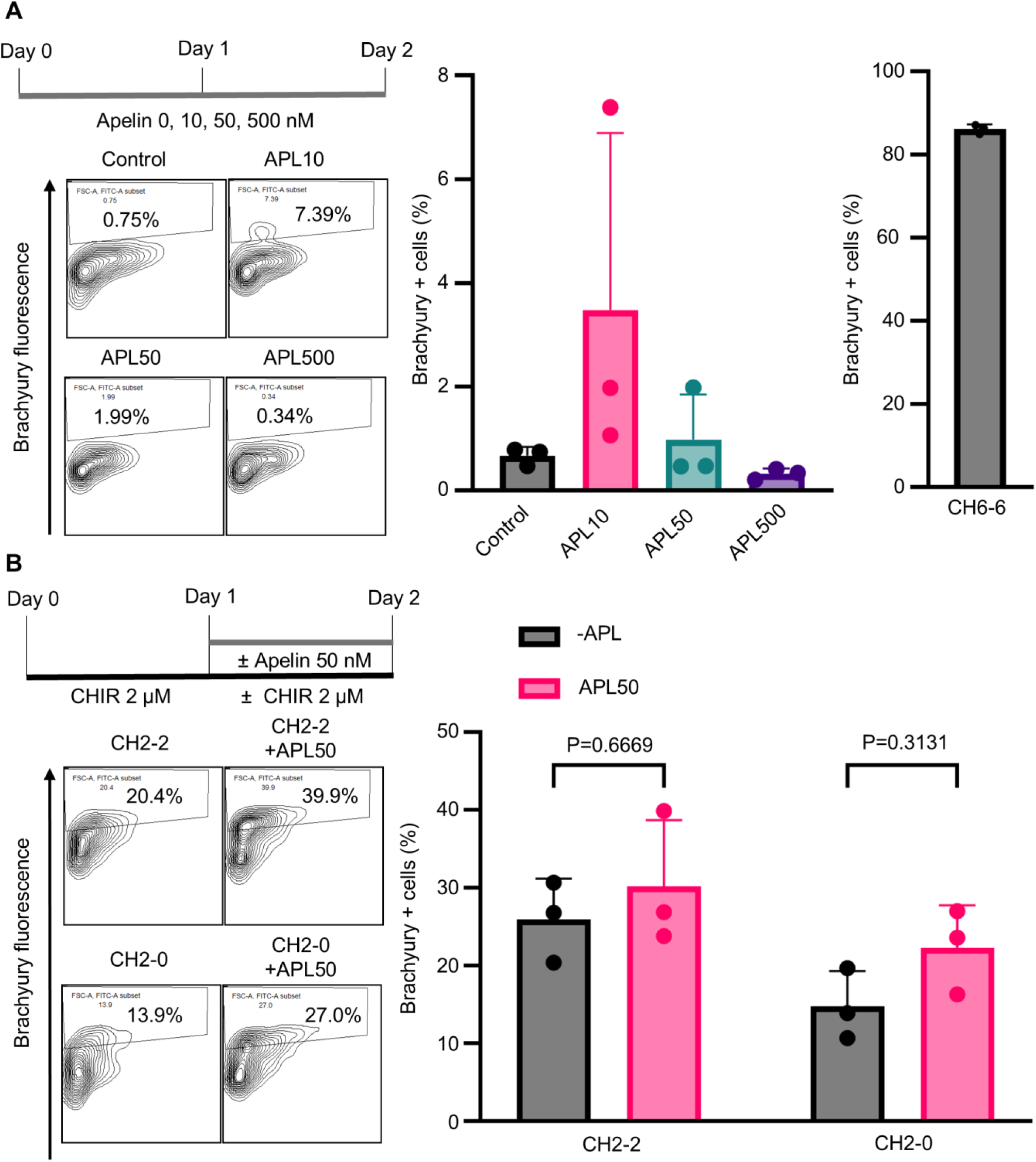
Apelin treatment did not significantly increase Brachyury expression during early hPSC differentiation. (A) Treatment schedule, representative flow cytometry plots and Brachyury-positive fractions after H9 cells received 0, 10, 50 or 500 nM [Pyr1]-Apelin-13 on days 0 and 1. The positive control received 6 µM CHIR99021 on both days (CH6-6). (B) Treatment schedules, representative flow cytometry plots and Brachyury-positive fractions after 2 µM CHIR99021 treatment with or without 50 nM Apelin on day 1. CH2-2 denotes CHIR99021 on days 0 and 1; CH2-0 denotes CHIR99021 on day 0 only. Brachyury expression was measured on day 2 in both experiments. Percentages in the contour plots indicate Brachyury-positive cells in the displayed wells. Bars show mean ± SD, with individual replicates plotted (n = 3 per condition). Apelin-only groups and the untreated control were compared by one-way ANOVA with Tukey’s multiple-comparisons test (overall P = 0.1952). The combination experiment used two-way ANOVA with Šídák-adjusted comparisons of Apelin treatment within each CHIR99021 schedule; adjusted P values are shown in B. APL, Apelin.

We next asked whether Apelin could enhance differentiation following Wnt pathway activation. H9 cells received a lower CHIR99021 concentration of 2 µM, either on both days 0 and 1 or on day 0 alone, with or without 50 nM Apelin on day 1. Apelin supplementation resulted in modest increases in the mean Brachyury-positive fraction under both CHIR99021 treatment schedules (Fig. 3B), but neither difference was statistically significant (Šídák-adjusted P=0.6669 and P=0.3131, respectively). Thus, Apelin did not significantly increase Brachyury expression under the tested conditions. The interactive analysis nonetheless connected a public dataset to a specific, experimentally testable hypothesis, while the experimental outcome showed why transcriptional enrichment alone is insufficient to predict the effect of adding a differentiation factor.

### 2.2 Autonomous single-cell analysis identifies reproducible candidates but depends on how the biological question is defined

We next asked whether a single prompt could direct an AI agent through an entire single-cell RNA-sequencing study, from finding and downloading suitable public data to analyzing the cells and proposing biological candidates. We focused on cell-surface antigens that might allow engineered immune cells to recognize tumor cells, building on our previous research in CAR-based immunotherapy [18,24–27]. Pancreatic ductal adenocarcinoma (PDAC) provided a solid-tumor setting in which candidate expression had to be distinguished from expression in normal pancreatic and stromal cells. Multiple myeloma (MM) provided a complementary setting in which the resulting candidates could be compared with established therapeutic antigens. For each disease, we initiated three separate runs with the same biological objectives and asked the agent to select datasets, establish its computational environment, perform the analysis and evaluate its conclusions without routine assistance. Repeating the task allowed us to examine both its ability to carry out the work and the consistency of the resulting candidates.

All six runs acquired datasets, filtered low-quality cells and assigned cell identities, and none requested scientific or routine programming assistance. Their subsequent analytical choices nevertheless differed. The PDAC runs analyzed three, four and three underlying tumor collections and retained 179,441, 279,924 and 76,221 cells, respectively (Fig. 4A). Runs 1 and 3 used the dataset reported by Steele et al. to identify initial candidates, whereas Run 2 used a reprocessed version of those tumor data together with additional donor-pancreas samples [28,29]. The runs also selected different additional PDAC datasets and normal-tissue references. Repeating the prompt therefore tested whether independent attempts to choose and analyze public data would reach similar conclusions, with each run making its own decisions about suitable inputs and methods.

**Figure 4.**
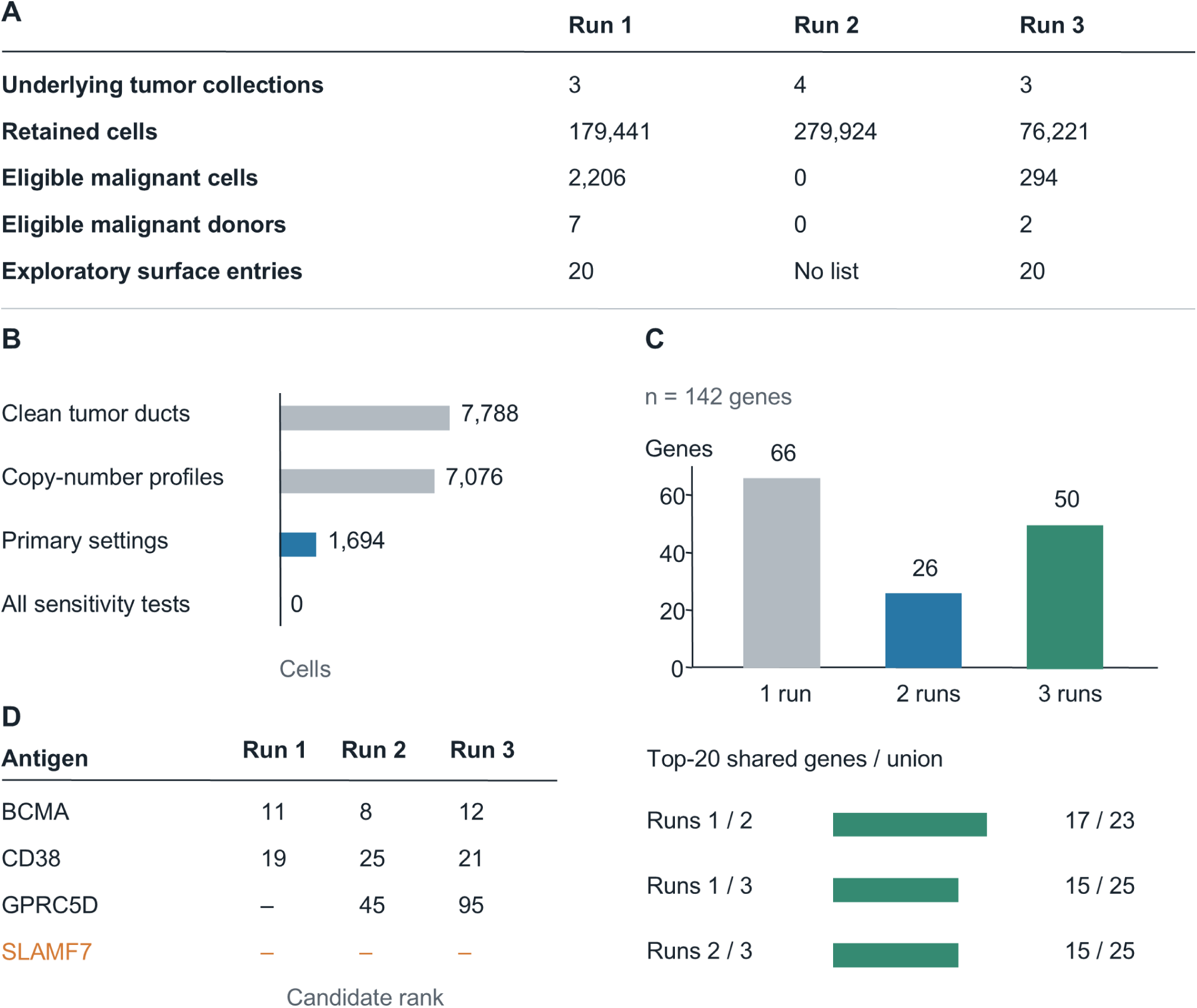
Repeated autonomous analyses of tumor-antigen candidates in pancreatic cancer and multiple myeloma. (A) Data coverage and results from three pancreatic ductal adenocarcinoma (PDAC) runs. Retained cells passed each run’s quality filters; eligible malignant cells additionally met its requirements for cancer-cell identification and the number of cells per donor. Runs 1 and 3 generated exploratory surface-gene lists, but no candidate met all target-selection criteria. Run 2 completed the analysis without a target list because its cancer-cell assignments were not stable when analysis settings were changed. (B) Successive stages of cancer-cell identification in Run 2. RNA expression was used to estimate large changes in chromosome copy number, which can help distinguish malignant from nonmalignant cells. The initial settings classified 1,694 cells as malignant, but no cells retained that classification across all required changes in copy-number window size and reference cells. (C) Number of multiple-myeloma (MM) candidates present in one, two or all three runs. The lists contained 86, 56 and 126 genes, comprising 142 different genes in total. (D) Ranks of four established MM antigens and agreement between the top-20 lists. BCMA is encoded by TNFRSF17; a dash indicates absence from the candidate list. Horizontal bars show the number of shared genes divided by the number of different genes in the two lists combined. Fifteen genes appeared in all three top-20 lists.

Identification of malignant epithelial cells was a major determinant of the PDAC results. Run 1 identified 2,206 eligible malignant cells from seven donors, but nonmalignant ductal cells from adjacent tissues were available for comparison from only three donors. Run 3 obtained 294 eligible malignant cells from two donors. Both runs retained 20 surface-associated genes for exploratory assessment, but the lists contained no shared genes, and all listed genes exceeded the respective expression limits in nonmalignant cells. In Run 3, six genes (GPRC5A, CEACAM6, LSR, F11R, DSG2 and CLDN4) met the predefined tumor-cell expression criteria in two additional primary-tumor datasets. However, their expression in nonmalignant cells prevented them from meeting the final target-selection criteria. The different lists therefore reflected differences in the malignant populations, surface-protein annotations and ranking rules, as well as the requirement to consider expression outside the tumor.

The second PDAC run completed a broader analysis that included six organ-donor pancreata, but it could not establish a stable set of malignant cells for target discovery. As in the other PDAC runs, Run 2 used RNA expression patterns to estimate changes in chromosome copy number, which are commonly observed in cancer cells and can help distinguish malignant from nonmalignant cells. It also tested whether the same cells were consistently identified as malignant when different analysis settings were applied. Of 7,788 tumor ductal cells that passed quality control, 7,076 had enough information for this assessment. The initial settings classified 1,694 cells from ten patients as malignant, but no cells were consistently classified as malignant across all tested settings, including changes in the size of the chromosomal regions examined and the reference cells used for comparison (Fig. 4B). The workflow therefore stopped before testing which genes were enriched in malignant cells and did not produce a target list. It nevertheless generated normal-reference expression summaries from 57,687 cells. The three PDAC runs consequently nominated no target for different reasons. The exploratory genes in Runs 1 and 3 were also expressed in nonmalignant cells, whereas Run 2 could not identify a sufficiently stable malignant-cell population for comparison.

The MM runs yielded more consistent candidates despite using different patient datasets. Run 1 used GSE193531 for candidate identification and GSE161801 for validation, while Run 2 reversed these roles. Run 3 used GSE232988 for candidate identification and GSE193531 for validation [30–32]. The three runs analyzed 10,754, 53,729 and 7,228 malignant plasma cells from 8, 19 and 22 donors, respectively. They identified 86, 56 and 126 candidate genes, representing 142 distinct genes overall. Among these, 50 genes were identified in all three runs, 26 in two runs and 66 in only one run (Fig. 4C; Supplementary Fig. S2 and Supplementary Table S3). Fifteen genes appeared among the top 20 candidates in all three runs. Comparisons between pairs of runs showed that 17, 15 and 15 of their top 20 candidates overlapped, corresponding to overlap ratios of 0.739, 0.600 and 0.600, respectively. When evaluated in separate MM datasets, 72, 42 and 55 candidates from Runs 1, 2 and 3 met their respective expression criteria, with 39 genes meeting these criteria in all three runs. Overall, many expression-based candidates were consistently identified across runs, although some of this agreement may reflect the reuse of patient datasets.

Comparison with established MM antigens clarified the biological meaning of this agreement. BCMA, encoded by TNFRSF17, ranked 11th, 8th and 12th, while CD38 ranked 19th, 25th and 21st (Fig. 4D and Supplementary Table S4). Both met the corresponding expression criteria in the separate datasets used for validation. GPRC5D appeared in two lists but did not meet those criteria in either validation analysis. SLAMF7 was absent from all three lists despite sufficient RNA expression because its database description identified it only as a membrane protein and did not satisfy the automated cell-surface annotation rule. Overall, the runs recovered two, three and three of the four reference antigens. The highest-ranked genes also included broadly expressed HLA and membrane-associated transcripts, because the MM ranking favored genes expressed in many tumor cells and many patients without requiring tumor-specific expression. Autonomous analysis therefore repeatedly recovered established antigens and a common set of expressed genes, while selecting targets for engineered immune cells still required assessment of cell-surface localization and expression outside the tumor.

### 2.3 Re-exploration without the source publication identified additional biological relationships in lung single-cell data

Single-cell studies generate information across many cell populations, although individual publications often focus on selected biological questions. We therefore asked whether AI could identify additional gene expression patterns and associations across cell populations in an existing dataset without being provided with information about the original study or its findings. This experiment used a human lung dataset containing epithelial, immune, vascular and stromal populations from patients with idiopathic pulmonary fibrosis (IPF) and nonfibrotic controls [33]. This multicellular dataset allowed us to examine both disease-associated changes within cell types and relationships between cell types across patients. The blind analysis received anonymized expression data and technical metadata, whereas a separate context-informed analysis was additionally told the tissue and disease labels. Both were instructed to survey the data broadly, including patterns beyond the principal difference between groups, and to complete their analyses before accessing the source publication.

The input comprised 89,326 cells from 12 patients with IPF and 10 controls, represented by 31 sequencing libraries. Without the tissue or diagnosis labels, the blind analysis identified alveolar and airway epithelial populations together with immune, endothelial and stromal cells, inferred a lung origin, and identified the IPF group as showing greater tissue remodeling. Both analyses recovered epithelial remodeling and changes in macrophage states, including greater representation of cells expressing SPP1 and GPNMB, while also identifying metabolic and stress-related gene-expression patterns. The two independently generated cell maps retained 87,705 and 87,698 cells after their respective quality-control steps. These findings provided a recognizable biological framework for examining additional questions rather than restricting the analysis to a single expected disease process (Fig. 5).

**Figure 5.**
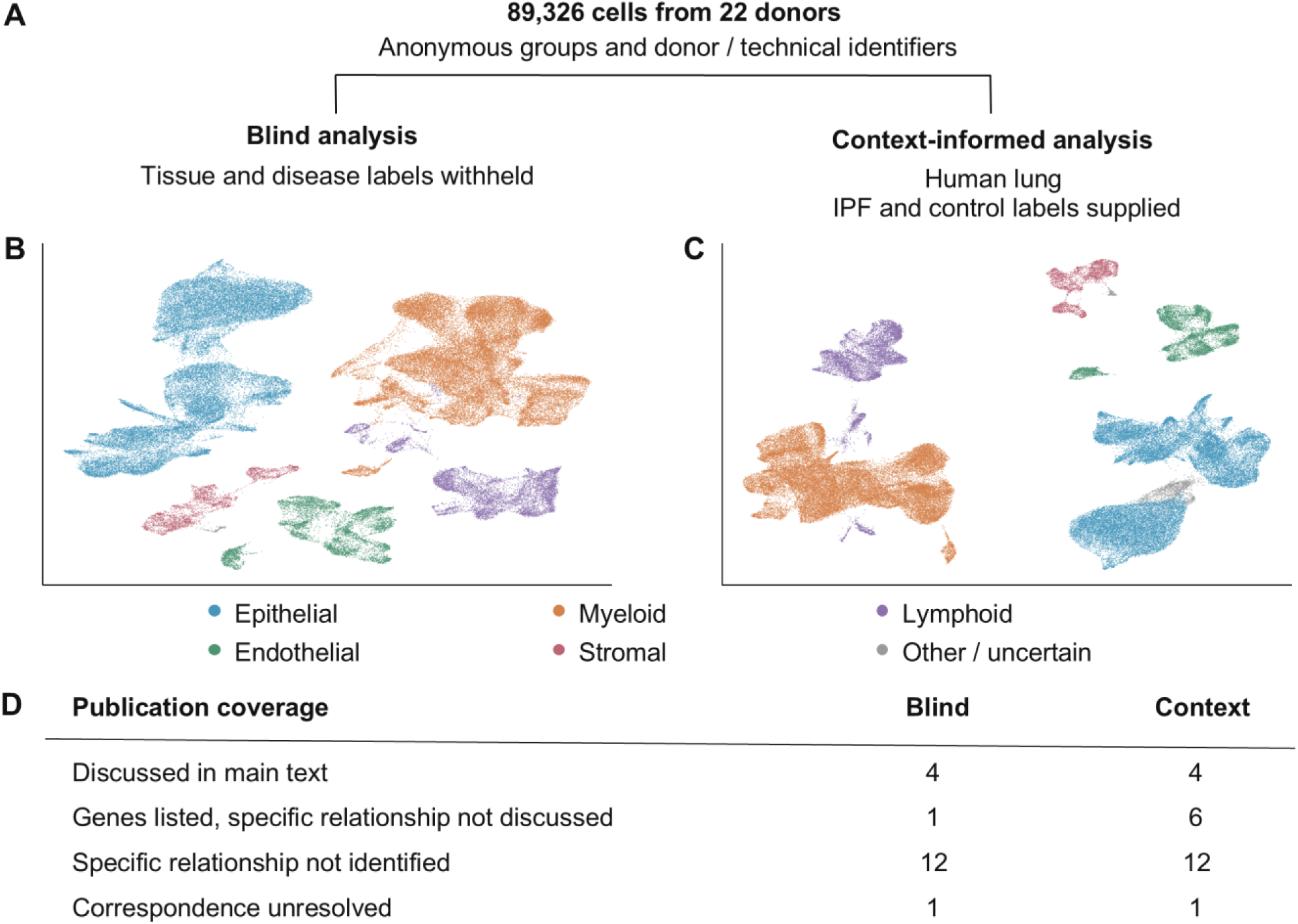
Blinded and context-informed reanalysis of a public lung single-cell dataset. **(A)** Study design. Both analyses received the same expression data and anonymized donor, group and technical identifiers. Only the context-informed analysis received the tissue and disease labels. The blind analysis used a separate account with personalization and memory disabled and a reset Work session. Both analyses were completed before comparison with the source publication. **(B,C)** Two-dimensional UMAP maps of 87,705 cells retained by the blind analysis and 87,698 cells retained by the context-informed analysis. Each dot is a cell, with similar expression profiles generally placed nearby. Colors denote broad cell groups. These independently calculated maps represent gene-expression similarity, not anatomical locations. **(D)** Comparison of 18 blind-analysis findings and 23 context-informed findings with the original publication. “Genes listed” indicates that relevant genes appeared in published tables, but the specific relationship proposed by the AI was not discussed. “Specific relationship not identified” indicates that the comparison did not locate that relationship in the publication. “Correspondence unresolved” indicates that a reliable match could not be established.

After both analyses were completed, a separately prompted AI comparison evaluated their findings against the original publication and its supplementary analyses. Four findings from each analysis corresponded to observations discussed in the main paper, while another 12 findings from each analysis described specific gene expression patterns or associations that were not identified in the publication. Some of the relevant gene expression results were already available in the original publication’s supplementary tables, but the AI analyses examined additional associations and expression patterns that had not been explicitly investigated. To assess whether these findings could be reproduced in other patients, a separately prompted AI analysis selected 13 questions for testing in a separate dataset, including seven from the blind analysis and six from the context-informed analysis (Supplementary Tables S5 and S6).

The blind analysis identified an association between antioxidant-related gene expression in alveolar type 2 (AT2) cells and secretory epithelial cells. An antioxidant-related expression score was calculated for each cell type in each patient using selected genes, including NQO1, HMOX1 and glutathione-related genes. Patients with higher expression scores in AT2 cells also tended to have higher scores in secretory cells, even after accounting for IPF status and specimen source (adjusted r=0.833, n=17 donors). The analysis also identified associations involving fatty-acid-oxidation-related genes in endothelial cells and macrophages, interferon-response genes in myeloid populations, and a small epithelial population expressing several neuroendocrine markers. The context-informed analysis generated additional questions involving increased CRYM expression in fibroblasts from IPF patients, higher glycolysis-related gene expression in endothelial cells, and differences in DNA-damage-response-related expression in AT2 cells and macrophages. Together, these findings provided specific gene expression patterns and associations for further evaluation beyond the principal IPF–control comparisons.

To examine whether these findings could be reproduced in other patients, a separately prompted AI analysis evaluated the 13 questions using another public lung dataset [34]. After excluding donors represented in the original study, the dataset included 28 patients with IPF and 42 nonfibrotic controls across 91 sequencing libraries. The number of donors included in each comparison varied according to the availability of sufficient cells from the relevant populations. Eight of the 13 questions showed the predicted expression differences or associations with statistical support. One additional question was supported by the detection of a predicted rare cell population, while three patterns were consistent with cellular stress or background RNA. The remaining question could not be evaluated because too few control cells of the required types were available (Fig. 6D). Detailed results and donor numbers are provided in Supplementary Table S6.

**Figure 6.**
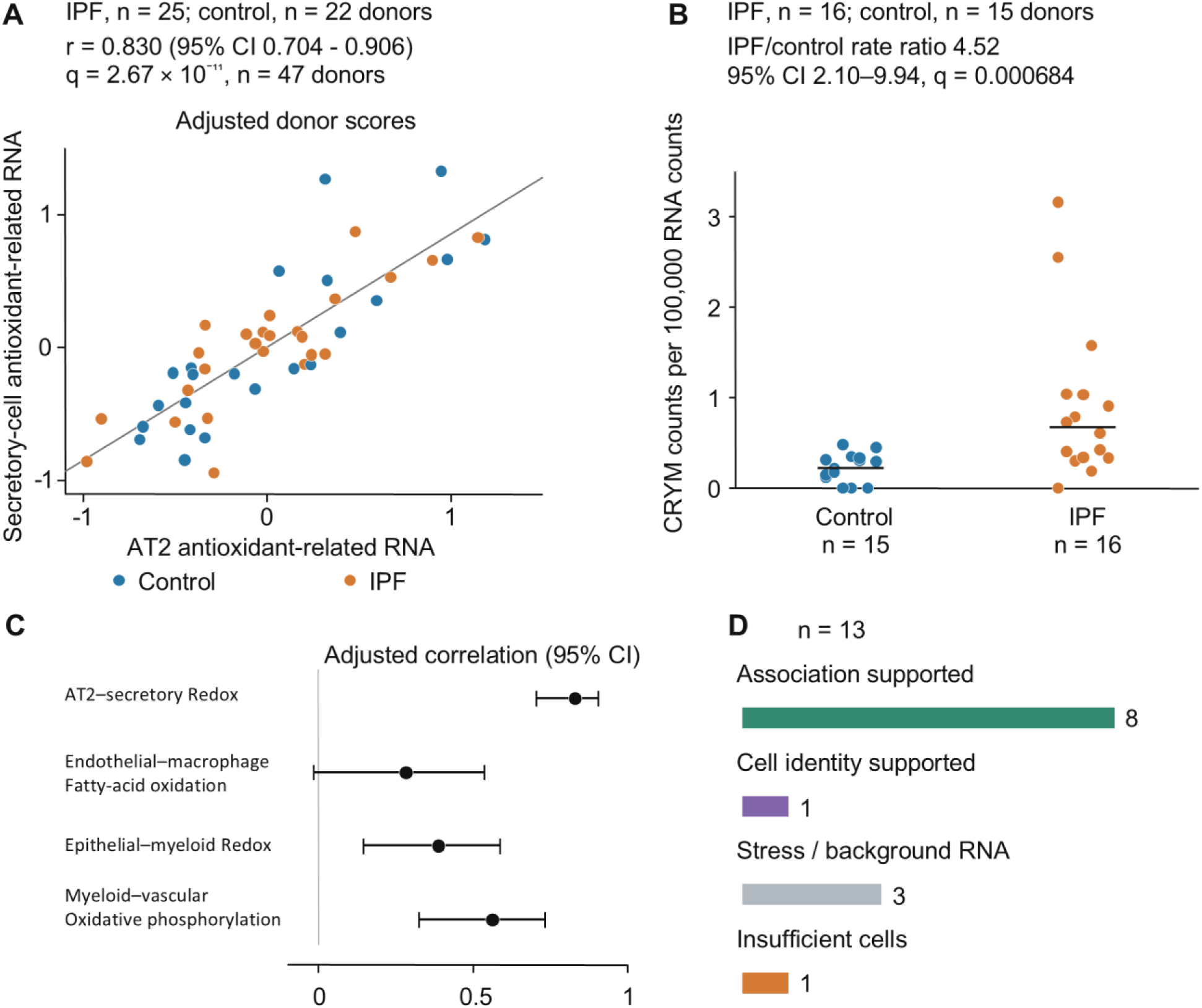
Testing additional lung expression patterns in other donors. The second dataset contained 28 IPF and 42 control donors after excluding donors represented in the original study. Individual tests used the subsets with enough cells of the relevant types. **(A)** Antioxidant-related gene expression in alveolar type 2 (AT2) and secretory epithelial cells from 47 donors (25 IPF and 22 controls). Each point represents one donor, colored by diagnosis. Scores summarize the predefined gene set in each cell population after adjustment for diagnosis, processing site, sequencing depth, mitochondrial RNA and library number. Positive and negative scores indicate expression above and below the level expected after adjustment, respectively; they are not transcript counts. The line shows a least-squares fit. **(B)** Fibroblast CRYM RNA in 16 IPF and 15 control donors, expressed as CRYM counts per 100,000 total fibroblast RNA counts. Points represent donors; horizontal lines show medians. The annotated IPF/control ratio and 95% CI are estimates from a negative-binomial count model. The joint q value summarizes evidence for both increased CRYM RNA and a larger fraction of CRYM-positive fibroblasts; smaller values indicate stronger statistical support. **(C)** Strength of four expression relationships between cell populations. Points show correlations after adjustment for diagnosis and technical measurements; bars show two-sided 95% CIs. Positive values indicate that the two expression scores increase together across donors; zero indicates no linear association. The endothelial– macrophage fatty-acid-oxidation result passed the prespecified one-sided test, although its two-sided CI spans zero and significance was lost with stricter cell filtering. **(D)** Outcomes of the 13 additional questions. Cell-identity support denotes recovery of a rare population, and stress/background denotes patterns whose biological interpretation remained unresolved. r, strength of the adjusted correlation; q, P value adjusted for multiple testing; CI, confidence interval describing uncertainty in the estimate; IPF, idiopathic pulmonary fibrosis. Redox refers to oxidation–reduction-related expression, and the pathway labels describe RNA patterns rather than measured metabolic activity.

One of the most consistent findings involved antioxidant-related gene expression in AT2 and secretory epithelial cells. The external analysis evaluated the same predefined gene set in both populations across 47 donors, including patients with IPF and nonfibrotic controls. As in the original dataset, higher expression scores in AT2 cells were associated with higher scores in secretory cells, even after accounting for IPF status and technical differences among samples (adjusted r=0.830, 95% confidence interval [CI] 0.704–0.906, q=2.67×10⁻¹¹; Fig. 6A). Each point in Figure 6A represents one donor. This result supports a reproducible association in RNA expression between the two epithelial populations rather than an increase or decrease specifically associated with IPF. Antioxidant activity itself was not measured.

Fibroblast CRYM expression also showed a disease-associated difference in the external dataset. Among 31 donors with sufficient fibroblasts, CRYM RNA expression relative to total fibroblast RNA was estimated to be 4.52 times higher in IPF patients than in nonfibrotic controls (95% CI 2.10–9.94; Fig. 6B). The proportion of fibroblasts expressing CRYM was also higher in IPF, with statistical support for both measurements (joint q=0.000684). Other supported findings included higher glycolysis-related gene expression in endothelial cells from IPF patients, MHC class II antigen-presentation-related gene expression in lymphoid cells, and coordinated antioxidant- and oxidative-phosphorylation-related expression across additional cell populations (Fig. 6C and Supplementary Table S6). These results provided further examples of disease-associated expression differences and cross-cell-population associations identified through AI reanalysis and supported in another patient dataset.

Not all findings were equally consistent. The association involving fatty-acid-oxidation-related genes in endothelial cells and macrophages met the initial statistical criterion (r=0.283, one-sided q=0.0376) but was no longer significant under stricter cell-quality filtering. The external analysis also detected neuroendocrine-like marker combinations in 78 epithelial cells from five donors, supporting the presence of the predicted rare population without establishing its enrichment in IPF. Other recurring patterns, including immunoglobulin RNA in unrelated cell types and shared heat-shock or erythroid RNA expression, could reflect technical factors rather than specific biological relationships. Overall, the blind and context-informed analyses generated complementary questions, several of which were supported in additional patients, while others remained sensitive to analysis settings or required alternative explanations (Supplementary Fig. S3 and Supplementary Table S6).

### 2.4 Integrating public molecular and functional data changed PDAC target prioritization

In our earlier PDAC and MM analyses, candidate tumor antigens were identified primarily based on their RNA expression in malignant cells. RNA abundance alone, however, does not establish whether the corresponding protein is expressed and accessible for therapeutic targeting, whether disrupting its function affects cancer-cell growth, or whether targeting it could also affect healthy tissues. To address these limitations, we conducted a separate AI analysis of PDAC therapeutic targets using additional types of publicly available data. We retained PDAC to connect this investigation to the earlier antigen screen and broadened the search to include intracellular proteins and extracellular factors as well as cell-surface antigens. The agent independently selected datasets and analytical methods, combining molecular measurements with functional screening data. Its instructions required selection and ranking criteria to be established before evaluating individual targets, followed by quantitative ranking before target-specific therapeutic literature review. Previous PDAC candidate lists were not supplied.

The agent selected complementary datasets addressing different aspects of target biology. RNA measurements from 60 paired tumors and adjacent tissues provided the initial screen, and a separate dataset of 118 tumors and 13 controls tested whether the expression differences recurred [35,36]. These bulk measurements describe whole tissue samples, so single-nucleus RNA sequencing of 108,964 nuclei from 18 untreated tumors was used to identify which cell populations expressed each candidate. Paired protein measurements then assessed whether increased RNA was accompanied by increased protein [37,38]. CRISPR screens tested whether disrupting a gene impaired cancer-cell growth, while normal-tissue RNA and protein references showed where the gene was expressed outside the tumor [39,40]. Additional genomic and regulatory measurements provided biological context. Together, these data addressed several requirements for a useful intervention beyond increased RNA expression in cancer tissue.

The combined screen identified 213 candidates through two routes: 207 genes had increased RNA expression in tumors, and six additional genes were selected because gene disruption affected PDAC cell growth more often than growth of other cancer cells. Among the 207 RNA candidates, 201 also showed increased expression in the separate dataset. Increased protein abundance was supported for 152 of all 213 candidates, and 149 had both confirmation of increased RNA in the separate dataset and protein support (Fig. 7A). CENPF, TOP2A and LAMA3 received the highest overall rankings (Fig. 7B and Supplementary Fig. S4A,B). Thus, many tumor-associated RNA changes were accompanied by increased protein, although measurements from whole tissue samples did not establish which cells produced that protein.

**Figure 7.**
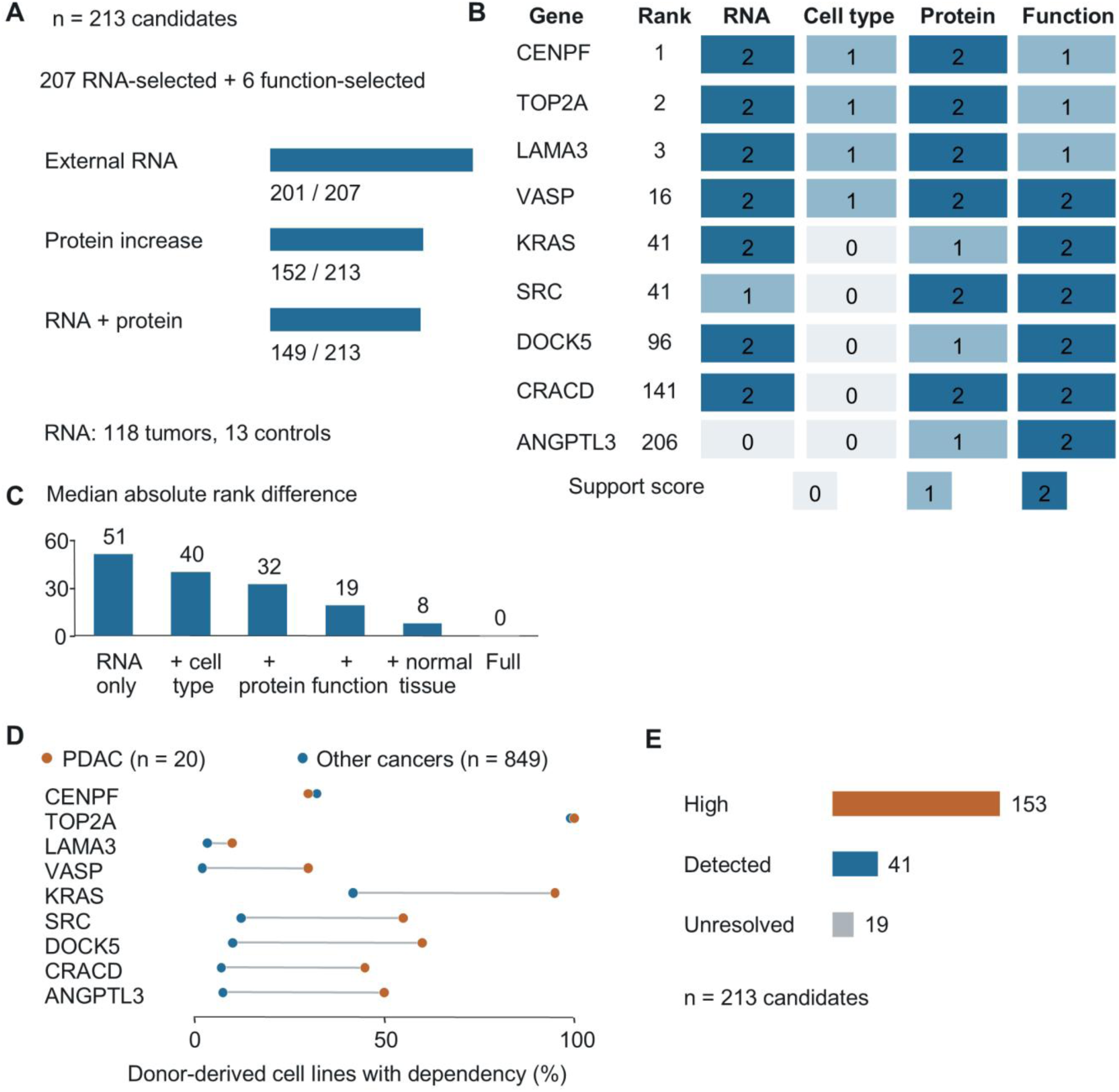
Combining public molecular and functional data changes the prioritization of PDAC candidates. (A) Support for 213 candidates: 207 selected through increased tumor RNA expression and six additional candidates selected through cancer-cell dependency. The bars show RNA confirmation in a separate study of 118 tumors and 13 controls, increased protein abundance in paired tumor and adjacent-tissue samples, and support from both RNA and protein measurements. RNA confirmation required a log2 expression difference ≥0.5 and adjusted P ≤ 0.05. Protein support required a mean paired increase ≥0.5 on the processed log scale, adjusted P ≤ 0.05 and at least 10 complete pairs. (B) RNA, cell-type, protein and functional evidence for the three highest-ranked candidates and the six additional candidates selected through dependency. Cell-type evidence describes expression in malignant versus other cell populations; functional evidence includes the effects of disrupting a gene in cancer cells. Scores range from 0 to 2, with higher values indicating stronger support under the criteria in the Supplementary Methods and Supplementary Table S8. (C) Changes in ranking as additional evidence types were added in sequence, using the same 213 candidates throughout. The graph shows the median absolute difference between each candidate’s rank in a reduced analysis and its rank in the full analysis; smaller values indicate closer agreement. Candidates tied under the ranking criteria shared a rank, so rank differences reflect both changes in support and the separation of previously tied candidates. The full analysis additionally incorporates regulatory associations and biological-function annotations. (D) Percentage of donor-derived cancer models classified as dependent on each gene, meaning that gene disruption reduced survival or growth in a CRISPR screen. Related cell lines were combined by donor, giving 20 PDAC and 849 other-cancer groups. (E) Normal-tissue expression among all 213 candidates. ‘High’ denotes ≥100 normalized transcripts per million (nTPM) or medium/high protein staining in a critical normal tissue; ‘detected’ denotes ≥10 nTPM without meeting the high-expression definition; ‘unresolved’ denotes insufficient reference information. These categories summarize expression rather than measured treatment toxicity. PDAC, pancreatic ductal adenocarcinoma.

CRISPR-based functional screening changed the interpretation of several candidates by revealing how gene disruption affected cancer-cell growth. When multiple cancer cell lines originated from the same donor, they were grouped together to avoid treating them as independent samples. TOP2A disruption impaired growth in all 20 PDAC donor groups, but the same dependency was present in 841 of 849 groups from other cancers. Its high RNA and protein expression therefore coincided with a requirement for growth shared across many cancers. In contrast, KRAS, DOCK5, SRC, ANGPTL3, CRACD and VASP were required for growth more often in PDAC than in other cancer types and entered the candidate list through this functional screen, although none met the RNA discovery threshold (Fig. 7D). KRAS, for example, was required in 19 of 20 PDAC groups compared with 355 of 849 other-cancer groups. Functional screening could therefore identify genes that would be missed by selecting only those with strongly increased tumor RNA expression. In a comparison with 18 genes associated with established or investigational treatment mechanisms, only KRAS occurred in the selected candidate set (Supplementary Table S9). Thus, the expression and dependency criteria also excluded several treatment-relevant mechanisms.

To assess the contribution of different evidence types, the analysis compared rankings of the same 213 candidates using RNA measurements alone and then successively adding cell-type information, protein measurements, functional data and normal-tissue expression. RNA measurements alone assigned the same highest score to 146 candidates, whereas additional evidence distinguished their levels of support. Using shared ranks for tied candidates, the median absolute difference from the full ranking was 51 with RNA evidence alone, 32 after adding cell-type and protein information, and 19 after adding functional data (Fig. 7C). These comparisons illustrate how additional measurements distinguished candidates that received similar support from RNA expression alone. The earlier single-cell screen used different datasets and focused specifically on cell-surface antigens for engineered immune cells, so its candidate lists were considered separately.

Normal-tissue measurements further clarified which candidates would require careful evaluation before intervention. High expression in normal tissues was detected for 153 candidates, lower but detectable expression for 41, and insufficient information was available for 19 (Fig. 7E). The resulting candidates therefore remain hypotheses for testing because the available measurements did not establish both protein expression in the intended tumor-cell population and a functional difference from healthy cells sufficient to support selective treatment. Nevertheless, combining the data made these hypotheses more informative by linking tumor RNA to protein abundance, identifying growth requirements missed by the RNA screen, and incorporating expression in normal tissues. A subset of 131 candidates had cell-surface or extracellular annotations and could be evaluated further where an accessible target is required.

As a separate demonstration, we asked the agent to create an interactive visualization using a publicly available pancreatic adenocarcinoma spatial transcriptomics dataset from 10x Genomics. The resulting viewer displayed 190,965 segmented cells at their measured tissue coordinates, with cell annotations grouped into six broad categories and ten selectable RNA expression overlays. Representative views of cell-type distribution and spatial expression of EPCAM and CTHRC1 are shown in Supplementary Fig. S5. Although similar visualization capabilities are available through existing platforms, this exercise demonstrated the ability of a general-purpose AI agent to create an interactive resource from public spatial data. The visualization was intended for data exploration and was not used for candidate prioritization in the multi-omics analysis [41]. The interactive viewer is available through the link provided in the Data and Code Availability section.

## 3. Discussion

This study examined the outcomes of using general-purpose AI to investigate public molecular data and develop questions within an experimental research project. Interactive guidance supported the transition from computational setup to a differentiation experiment, and autonomous prompts directed dataset acquisition, analysis and reporting across several disease settings. The outcomes varied with the question and the available evidence. The Apelin experiment did not establish the proposed differentiation effect, repeated MM analyses identified a common set of RNA candidates, blind and context-informed lung analyses generated additional expression patterns and associations that could be tested in other patients, and integration of additional data changed PDAC therapeutic target prioritization. These outcomes illustrate how AI can support different stages of biological research, while showing the importance of evaluating computational findings through additional analyses and experimental testing.

The Apelin investigation provides a direct example of the transition from an expression pattern to an experimental hypothesis. Increased APLNR expression in primitive-streak-like cells suggested that its ligand might influence differentiation. However, expression in an emerging cell state does not establish that receptor activation initiates that state. The analyzed gastruloid samples described cells after differentiation and after reseeding, rather than following the onset of mesendoderm formation. The resulting hypothesis therefore required an intervention experiment. Neither Apelin alone nor its addition to the tested low-dose CHIR99021 schedules produced a statistically significant increase in the Brachyury-positive fraction.

Prior Apelin studies help place this result in a developmental context. Yu et al. examined Apelin during cytokine-directed hematopoietic differentiation, and Wang et al. reported improved cardiac differentiation in embryoid-body-based protocols [20,21]. Macrae et al. linked APLNR signaling to the efficiency and function of hESC-derived cardiomyocytes [22]. Of particular relevance, Şişli et al. reported increased Brachyury expression after Apelin-13 treatment during CHIR-containing H9 differentiation [23], although their study used embryoid-body culture with different treatment and analysis time points. These studies provide functional precedent for investigating Apelin during human stem cell differentiation, but do not establish that the ligand is sufficient to initiate mesendoderm under the conditions tested here. Considering the starting cell state, concurrent signals and timing of the readout is therefore essential when translating a literature-supported pathway into an experimental treatment.

The repeated autonomous analyses examined whether an AI agent could complete a broad public-data investigation and identify similar biological candidates across separate runs. MM showed substantial candidate recurrence and repeated recovery of BCMA and CD38. At the same time, SLAMF7 was consistently excluded because the automated cell-surface annotation criteria did not recognize it as an eligible target. Thus, agreement between runs can reflect both reproducible expression patterns and limitations shared by the analytical methods. The PDAC analyses were particularly sensitive to how malignant epithelial cells were identified and which nonmalignant populations were used for comparison. Run 2 analyzed the largest cell collection, but its malignant-cell classifications were not consistent when the copy-number analysis settings were changed. These results show that the datasets, analytical methods and selection criteria chosen by an AI agent can substantially influence the resulting candidates, even when the same biological objectives are provided.

The contrasting outcomes between PDAC and MM may also reflect differences in their underlying biology. PDAC is characterized by substantial tumor heterogeneity and a complex stromal microenvironment, which can complicate the identification of malignant epithelial cells and the selection of antigens with limited expression in normal tissues [42,43]. In contrast, MM arises from plasma cells and has several well-established therapeutic antigens, including BCMA, which is targeted by clinically approved CAR T-cell therapies [44,45]. These differences may have contributed to the more consistent recovery of candidates in MM, although the effects of disease biology cannot be separated from differences in dataset composition and analytical criteria in the present study. Thus, the contrasting results should not be interpreted solely as differences in AI performance.

The lung investigation illustrates how AI analyses can examine additional questions within a published dataset. Published single-cell studies contain measurements across many cell populations, although individual publications often focus on selected biological questions. The blind and context-informed analyses examined gene expression patterns and associations across cell populations without being provided with the original publication or its findings. Several resulting questions involved information already available in the supplementary tables but examined relationships that had not been explicitly investigated in the original study. The association between antioxidant-related gene expression in AT2 and secretory epithelial cells and the increased CRYM expression in fibroblasts from IPF patients were supported in another patient dataset. These results suggest that AI-assisted reanalysis can identify additional questions within existing measurements without requiring that every observation represent an entirely new gene or biological pathway.

Testing these findings in other patients also helped assess their consistency and limitations. Some expression patterns remained supported under different analysis settings, whereas the association involving fatty-acid-oxidation-related genes in endothelial cells and macrophages was no longer statistically significant with stricter cell-quality filtering. Other recurring patterns were consistent with cellular stress or RNA originating from unrelated cells during sample preparation. Although donors represented in the original study were excluded, the additional patients came from a collection network shared with the original study. These results therefore provide evidence that several AI-identified expression patterns recur in other patients, while their biological significance remains to be established. The blind analysis also demonstrated that the agent could infer the tissue origin and disease-associated changes without being given the tissue or diagnosis labels. However, the agent’s general biological knowledge and the instructions in the prompt continued to influence its analysis, so the results should not be interpreted as completely independent biological discovery.

The broader PDAC analysis demonstrated how integrating different types of public data can change therapeutic target prioritization. Protein measurements provided additional evidence for tumor-associated RNA expression, while CRISPR-based functional screening identified genes whose disruption affected cancer-cell growth even when they did not meet the RNA expression criteria. Normal-tissue expression data also helped identify candidates that might present challenges for selective targeting. TOP2A illustrates how increased tumor expression and strong functional dependency do not necessarily indicate a dependency specific to PDAC, as its disruption also affected many other cancer types. In contrast, KRAS and five other genes entered the candidate list through functional screening despite failing the RNA discovery threshold. These comparisons illustrate how additional measurements can distinguish candidates that received similar support from RNA expression alone. However, these computational results do not establish whether targeting the selected proteins would selectively affect cancer cells while sparing healthy tissues, which requires further experimental evaluation.

These investigations complement specialized single-cell and biomedical-agent studies by examining several applications of general-purpose AI within an experimental research project. AI-assisted prompt development translated biological objectives into detailed analysis instructions, while the AI systems contributed dataset selection, computational methods, code execution and proposed interpretations. The results also showed that the value of AI-assisted analysis depends on how its findings are evaluated. Repeated analyses can assess candidate consistency, additional patient datasets can test whether expression patterns recur, and laboratory experiments can examine whether computational hypotheses produce the predicted biological effects. Together, these approaches demonstrate how general-purpose AI can support biological data analysis and hypothesis generation, while emphasizing the distinction between computational findings and experimentally established biological effects. These investigations were not intended as a systematic benchmark of AI systems, but rather as an examination of their practical use in addressing biological questions within an experimental research setting.

## 4. Materials and Methods

### 4.1 Study design and AI assisted workflows

The datasets are summarized in Supplementary Table S1. The study comprised interactive analysis of hPSC differentiation, repeated autonomous tumor-antigen discovery, blinded and context-informed re-exploration of lung single-cell data, and integration of multiple public-data types for PDAC target prioritization. Interactive analysis used Gemini Pro 3.1 through the Gemini conversational interface. The autonomous analyses used GPT-6 Astra with the Extra high or Ultra reasoning setting in ChatGPT Work, which provided a general-purpose agentic interface and a hosted computational environment. We defined the biological objectives and used AI assistance to develop detailed analysis instructions, which we reviewed and refined before submission. These instructions emphasized explicit biological comparisons, donor-level analysis where applicable, evaluation in additional datasets and separation of observations from proposed mechanisms. Within the instructions and data supplied for each autonomous task, the agent selected analytical procedures and settings, acquired additional public data where required, executed code, resolved routine software errors and generated results and biological interpretations. In the interactive workflow, we executed AI-provided code and performed the hPSC differentiation experiments. Available prompts, their provenance and run-specific software settings are provided in the supplementary materials and the computational repository.

For the repeated autonomous analyses, execution autonomy was assessed by whether scientific decisions or routine programming assistance were requested after the initial prompt. User-initiated pauses and resumptions were considered separately. Candidate consistency was assessed using the gene lists and rankings produced by each run. Comparisons between runs were initiated through additional prompts. In the lung study, separate prompts directed the initial blind and context-informed analyses, comparison with the source publication, and external testing. The publication comparison used the completed analyses together with the paper and its supplementary results. The AI agent independently selected the questions and established the statistical criteria for external testing before examining the external expression outcomes. Public data were analyzed at the biological-replicate level when testing differences or correlations across patients.

### 4.2 Interactive analysis of hPSC gastruloid scRNA-seq data

Gemini was instructed to guide dataset selection, software installation, scRNA-seq analysis and candidate prioritization through successive exchanges. We executed the supplied R commands and returned the outputs before proceeding to the next step. The analysis used GSE144897, comprising two H1 gastruloid replicates collected after 44 h of BMP4 treatment and cells profiled 48 h after dissociation and reseeding [19]. Data were analyzed in R 4.6.1 using Seurat 5.5.1. Cells expressing more than 1,000 and fewer than 8,000 genes, with less than 10% mitochondrial transcripts, were retained. Counts were normalized to 10,000 per cell and log-transformed. The 2,000 most variable genes were scaled and used for principal-component analysis. The first 20 principal components were used for nearest-neighbor clustering at a resolution of 0.5 and UMAP visualization.

Cell-state annotation used POU5F1, T, MESP1 and SOX17 expression. Clusters 0, 6 and 7 were grouped as putative pluripotent cells, cluster 1 as primitive-streak-like cells, and clusters 3 and 9 as mesoderm. Differential expression between the primitive-streak-like and putative pluripotent populations used Seurat FindMarkers with only.pos=TRUE and min.pct=0.25. Candidate genes were ranked by average log2 fold change. APLNR, APLN, APELA, GRP and GRPR expression was examined to prioritize experimentally accessible signaling candidates.

### 4.3 hPSC culture and differentiation

H9 human embryonic stem cells (WiCell) were maintained on Matrigel-coated plates in mTeSR Plus medium (STEMCELL Technologies, 100-0276) at 37°C and 5% CO2, with daily medium replacement. Plates were coated with 0.08 mg/mL Matrigel (Corning, 354230) in DMEM/F12 for at least 2 h at 37°C. Cells at 70–80% confluence were dissociated with 1 mM EDTA and passaged at 1:10 in medium containing 5 µM Y-27632 (Cayman Chemical, 10005583).

Differentiation was initiated at 70–80% confluence by replacing the maintenance medium with RPMI 1640 supplemented with B27 minus insulin. Cells received [Pyr1]-Apelin-13 (MedChemExpress, HY-P1033) at 10, 50 or 500 nM on days 0 and 1. Untreated cells and cells receiving 6 µM CHIR99021 (Cayman Chemical, 13122) on both days served as controls. In the combination experiment, cells received 2 µM CHIR99021 on days 0 and 1 or on day 0 alone, with or without 50 nM Apelin on day 1. Medium was replaced on day 1, and Brachyury expression was measured on day 2.

### 4.4 Flow cytometry and statistical analysis

Cells were dissociated with 0.05% trypsin-EDTA, fixed in 1% paraformaldehyde for 20 min at room temperature, and permeabilized in 90% methanol overnight at −20°C. After washing in PBS containing 2.5% BSA and 0.1% Triton X-100, cells were incubated with anti-Brachyury antibody (R&D Systems, AF2085, 1:100) overnight at 4°C, followed by chicken anti-goat IgG– Alexa Fluor 488 (Invitrogen, A-21467, 1:1000). Data were acquired on a BD Accuri C6 Plus and analyzed using FlowJo 10.8.1. Brachyury-positive fractions were calculated within the forward- and side-scatter cell gate.

Data are presented as mean±SD of three replicates per condition. Statistical analysis used GraphPad Prism 11. Apelin-only conditions and untreated controls were compared by ordinary one-way ANOVA with Tukey’s multiple-comparisons test. The CHIR99021–Apelin experiment used ordinary two-way ANOVA, with Šídák-adjusted comparisons of Apelin treatment within each CHIR99021 schedule. P<0.05 was considered statistically significant.

### 4.5 Autonomous tumor-antigen discovery and comparison of repeated runs

We initiated three separate autonomous analyses for PDAC and three for MM using disease-specific prompts with common instructions. Each prompt specified the biological objective, use of public human single-cell RNA-sequencing data, donor-level analysis, separate datasets for identifying and testing candidates where feasible, and assessment of candidate expression in nonmalignant cells. The primary objective was to identify cell-surface antigens for engineered immune-cell targeting, while intracellular genes and pathways were considered secondary objectives. The agent selected datasets and software, installed dependencies, performed the analyses and addressed routine computational errors. Within the prompt’s requirements, the agent established candidate-selection rules before examining target results and fixed the candidate lists before candidate-specific literature review or clinical benchmarking. The prompts and detailed computational procedures are provided in the Supplementary Methods.

Analyses used Python and Scanpy in hosted Linux environments. Quality control considered detected genes, library size, mitochondrial RNA and potential doublets separately for each dataset. Counts were normalized to 10,000 per cell and log-transformed for dimensionality reduction and marker assessment. Principal-component analysis, neighbor graphs, Leiden clustering and UMAP supported cell annotation. Multi-gene lineage markers and sample metadata informed cell identities. The PDAC analyses additionally used RNA-derived copy-number patterns to distinguish malignant epithelial cells from nonmalignant ductal cells. MM analyses combined plasma-cell expression programs with available neoplastic or clonal annotations. Donor identifiers were retained throughout, and technical samples from the same donor were combined for donor-level summaries. Dataset allocation, filtering thresholds, malignancy definitions and ranking functions were selected independently within each run and are reported in Supplementary Table S2.

For PDAC, Runs 1 and 3 used GSE155698 for discovery, with additional tumor datasets and normal references selected by each run. Run 2 used GSE229413, which combines reprocessed Steele tumor data with donor-pancreas samples, and analyzed GSE212966, GSE205013 and GSE154778 as additional collections [28,29,46–48]. The original and reprocessed Steele tumor data were treated as the same biological collection. Surface-associated genes were assessed for expression among malignant cells and across patients, with nonmalignant pancreatic and stromal populations used to assess specificity. In Run 2, malignant calls also had to remain supported when copy-number windows and reference populations were varied. This requirement was not met, so no malignant-versus-normal gene-expression model was fitted in that run.

For MM, Runs 1 and 2 exchanged GSE193531 and GSE161801 as datasets for identifying and testing candidates; Run 3 used GSE232988 for discovery and GSE193531 for validation [30–32]. Surface annotation was based on UniProt localization and topology. Candidate selection combined the fraction of expressing malignant cells, the proportion of positive patients and expression abundance. Candidate expression was then evaluated in the separate dataset used for validation using the original run-specific thresholds. After candidate selection, each workflow constructed a reference set of established MM antigens from regulatory and clinical sources. BCMA, CD38, GPRC5D and SLAMF7 formed the common four-antigen denominator.

Separately prompted comparison analyses assessed dataset selection, candidate membership and rank across runs. Gene identities were matched using the recorded gene symbols or Ensembl identifiers, and top-5, top-10 and top-20 overlaps were calculated as the number of shared genes divided by the number of distinct genes in the compared lists. Comparisons involving a run without a candidate ranking were recorded as unavailable. Candidate recurrence and recovery of known MM antigens were analyzed separately from expression confirmation in another patient dataset.

### 4.6 Blinded and context-informed analysis of lung single-cell RNA-seq data

We used the IPF and nonfibrotic-control subset of GSE135893 [33], comprising 89,326 cells from 22 donors and 31 sequencing libraries. Before the blind analysis, disease groups were renamed Group A and Group B, and donor, library, specimen-source, processing-site, flowcell and chemistry identifiers were replaced with anonymous codes. Repository accessions, original sample names, cell annotations and other identifying study information were excluded. We initiated the blind run using a separate account with personalization and memory disabled after resetting Work. Its instructions prohibited retrieval of previous conversations or analyses, identification of the source study, and disease-specific literature searches. The analysis was allowed to use general biological knowledge and software documentation. The context-informed run received the same anonymized data together with the human lung, IPF and control labels, but not the source publication, original annotations or blind-analysis results. Each analysis was finalized before comparison with the publication.

Both prompts requested autonomous quality control, cell-type annotation, donor-level comparisons, broad analysis of groups of genes associated with biological processes and investigation of relationships across cell populations. The AI selected computational parameters independently in each run. Primary filtering required at least 1,000 detected genes and less than 20% mitochondrial RNA; the context-informed analysis also excluded extreme library-wise count outliers. The blind analysis used log-normalized counts per 10,000, 3,500 variable genes, 50 principal components and a 20-neighbor graph based on 40 components. The context-informed analysis used 2,500 variable genes, 40 principal components and 20 neighbors. Both used Leiden clustering, UMAP and multiple markers for cell annotation.

For gene-expression comparisons, counts were summed within each donor and cell population, with donors treated as biological replicates. Models included disease group and processing site, with separate assessments of specimen source, quality control, normalization and individual-donor influence. Eligibility required at least 20 cells per donor and five donors per group in the blind analysis or four per group in the context-informed analysis. The Benjamini–Hochberg procedure controlled the false discovery rate within the specified test families. Full analytical settings and sensitivity analyses are described in Supplementary Methods.

### 4.7 Publication comparison and testing in external patients

After completion of the blind and context-informed analyses, a separately prompted AI comparison assessed their findings against the original paper and supplementary results, distinguishing relationships discussed in the publication from gene-level information listed without interpretation of the same relationship. A subsequent AI analysis evaluated 13 additional questions in GSE227136 [34]. Donors matching any of the 30 donors in the full original study were excluded, yielding 70 external donors. The AI agent independently defined the cell populations, gene panels, predicted directions and statistical criteria before evaluating the external expression outcomes. Two myeloid-state comparisons and three additional technical controls were analyzed separately.

External analyses used expression summaries for each donor and cell population, retaining only donors with enough cells for the specified comparison. To summarize a biological process, the external analysis combined the standardized expression of a predefined set of related genes into one score for each donor and cell population. Correlations tested whether these scores varied together in different cell types from the same donors, after accounting for diagnosis, processing site, sequencing depth, mitochondrial RNA and library number. CRYM was evaluated using a negative-binomial count model that accounted for the total fibroblast RNA measured in each donor, together with a test of the fraction of fibroblasts expressing CRYM. Directional tests were corrected for multiple comparisons within predefined families. Additional analyses assessed whether conclusions changed with stricter cell filtering, alternative cell definitions or removal of individual donors. Rare-cell identity and recurring background-RNA patterns were evaluated separately from disease-associated expression changes.

### 4.8 Public-data integration for PDAC target prioritization

The autonomous prompt specified PDAC therapeutic target discovery without restricting the data modality or target class. The agent selected datasets, assigned discovery and validation roles, and defined the scoring criteria before target prioritization. Target-specific therapeutic literature was reviewed after completing the quantitative ranking. Previous single-cell target lists were not supplied.

Processed microarray expression in GSE62452 was analyzed using paired t tests for 60 tumors and matched adjacent tissues [35]. Probes with unambiguous gene mappings were averaged within each sample. RNA candidate selection required a mean log2 expression difference ≥1 and Benjamini–Hochberg adjusted P≤0.05. Replication in GSE62165, comprising 118 tumors and 13 controls, used Welch’s t tests with a log2 difference ≥0.5 and adjusted P≤0.05 [36]. GSE71729 provided additional transcriptomic comparisons [49].

Cell-type expression was assessed in 108,964 nuclei from 18 untreated GSE202051 tumors using the published annotations [37]. Counts were aggregated by donor and cell population, requiring ≥20 nuclei per population and ≥5 paired donors for comparisons. CPTAC proteomic data included up to 62 tumor–adjacent pairs [38]. Protein support required a mean paired change ≥0.5 on the processed log scale, adjusted P≤0.05 and ≥10 complete pairs, without imputation. Assays from the same CPTAC patients were treated as complementary measurements from the same patients.

Functional screening used Project Score binary BAGEL2 dependency calls for 17,645 genes [39,40]. Related cell-line models were grouped by donor, yielding 20 PDAC and 849 nonpancreatic cancer units. Selective dependency required a prevalence difference ≥0.20 and a one-sided Fisher’s exact test adjusted P≤0.05. Normal-tissue expression was assessed using Human Protein Atlas RNA and immunohistochemistry data, with GTEx as an alternative RNA reference. Expression was classified as high at ≥100 nTPM or medium/high protein staining in a critical normal tissue, and present at ≥10 nTPM without a high-expression designation.

The union of RNA and functional candidates was scored across ten biological evidence categories, each receiving 0–2 points where measured. Normal-tissue expression and therapeutic accessibility were considered separately from the support score. Candidates were ranked by total support, support across separate patient datasets and conflicting evidence within each advancement category. Candidates tied under the ranking criteria shared the same rank. Alphabetical ordering was used only to select and display a fixed number of candidates from tied groups. Recalculating the rankings with successively added evidence types used the same 213 candidates throughout. Supplementary Table S8 summarizes the evidence categories and candidate-level results, and Supplementary Table S9 reports rankings obtained with different combinations of evidence. The complete component scores and scoring code are available in the computational repository.

### 4.9 Spatial visualization

A public 10x Genomics Xenium pancreatic adenocarcinoma dataset was visualized using measured cell-centroid coordinates and provider annotations [41]. All 190,965 cells were retained. Ten RNA overlays displayed assigned transcript counts, and the provider labels were grouped into six broad cell families for navigation. The two-dimensional coordinates were preserved without assigning anatomical depth. Dataset attribution and the CC BY 4.0 license were retained. Additional details are provided in the Supplementary Information.

## 5. Data and code availability

Public datasets are identified by repository accession and primary publication in the Methods and supplementary materials. Computational records supporting this study, including available prompts and execution records, analysis code, result tables, selected processed data and computational figure materials, are available through Zenodo at https://doi.org/10.5281/zenodo.23290610. The deposit includes an indexed file list and distinguishes original run-associated prompts from later supplied or reconstructed copies. The interactive pancreatic spatial transcriptomics viewer is publicly available at https://scintillating-jin-8e1ba7.netlify.app/model-02.

## AI assisted language editing

AI assistance was used to revise the language of the manuscript.

