## Supplementary information for "General Purpose AI for Public Data Analysis and Experimental Hypothesis Generation"

### Supplementary Methods

#### S1. Analysis instructions and computational workflows

We developed the analysis instructions from the biological question for each part, using AI assistance to expand the initial goals into detailed computational tasks. We reviewed and refined these instructions before execution. Part 1 used an interactive workflow in which Gemini Pro 3.1 provided instructions and R code, and we executed the code and returned outputs or errors. Parts 2–4 and the spatial companion used GPT-6 Astra with the Extra high or Ultra reasoning setting in ChatGPT Work. These agentic workflows could obtain public data, write and execute code, and prepare results within the same task. The instructions asked the agent to document its analytical choices and retain the numerical outputs underlying its conclusions. The purposes of the available analysis instructions are summarized in the Supplementary Prompt Summary.

The instructions addressed different questions. Part 1 sought experimentally testable signals for human pluripotent stem cell (hPSC) mesendoderm differentiation. Part 2 sought tumor antigens that might be targeted by engineered immune cells and tested whether separate autonomous runs produced similar results. Part 3 examined whether an analysis performed without knowledge of the source publication would identify additional relationships within a public dataset. Part 4 allowed the use of diverse public molecular and functional data to investigate pancreatic cancer targets beyond the cell-surface and single-cell RNA focus of Part 2. The pancreatic spatial viewer was developed as a separate resource for exploring cell locations and selected gene-expression patterns in tissue.

For the autonomous analyses, the agent selected datasets and analytical parameters within the requirements of the submitted prompt. Expression inference used donors or specimens as the statistical units where these identifiers were available. Cell-level summaries described the composition or expression distribution within samples. Repeated computational runs were compared using their resulting candidate lists and rankings.

#### S2. Interactive analysis of gastruloid single-cell data

Using the interactive AI-guided workflow, we analyzed GSE144897, a single-cell dataset from micropatterned H1 human embryonic stem cell gastruloids [1]. Two samples were collected after 44 h of BMP4 treatment, each comprising 36 pooled gastruloids. A third sample was collected 48 h after cells had been dissociated and reseeded. The samples therefore represented differentiated cell states and a reseeding condition. The interactive analysis used R 4.6.1 and Seurat 5.5.1.

The three count matrices were imported separately, retaining cells with at least 200 detected genes, and merged after assigning sample-specific barcode prefixes. The merged object contained 3,396 cells. We retained cells with more than 1,000 and fewer than 8,000 detected genes and less than 10% mitochondrial RNA, leaving 3,311 cells. Counts were normalized to 10,000 per cell and log transformed. The 2,000 most variable genes were selected using the variance-stabilizing transformation method, followed by scaling and principal-component analysis. The first 20 principal components were used to construct the neighbor graph and UMAP; clustering used a resolution of 0.5.

We examined POU5F1, T, MESP1 and SOX17 expression to identify relevant cell states. T was the gene symbol used for TBXT in the input annotation. Clusters 0, 6 and 7 were grouped as putative pluripotent cells, cluster 1 as primitive-streak-like cells, and clusters 3 and 9 as mesoderm-like cells. These groups contained 1,023, 562 and 496 cells, respectively. Marker analysis compared the primitive-streak-like and putative pluripotent groups using FindMarkers with positive markers only and a minimum detection fraction of 0.25. The 25 genes with the largest average log2 fold changes were retained. APLNR, APLN, APELA, GRP and GRPR were subsequently visualized across the selected groups. The retained UMAP describes relationships among observed cell states; it was not fitted as a developmental time course.

**S3. hPSC culture, Apelin treatment and Brachyury analysis**

H9 human embryonic stem cells (WiCell) were maintained in mTeSR Plus medium (STEMCELL Technologies, 100-0276) on Matrigel-coated plates at 37°C and 5% $CO_2$, with daily medium replacement. Matrigel (Corning, 354230) was diluted to 0.08 mg/mL in DMEM/F12 (Gibco, 11330-032) and incubated on plates at 37°C for at least 2 h. At approximately 70–80% confluence, cells were washed with calcium- and magnesium-free PBS (Corning, 21040CV), dissociated with 1 mM EDTA (Invitrogen, 15575-038), and passaged at a 1:10 ratio in medium containing 5 µM Y-27632 (Cayman Chemical, 10005583).

Differentiation was initiated at approximately 70–80% confluence by replacing the maintenance medium with RPMI 1640 (Gibco, 11875093) supplemented with B27 minus insulin (Gibco, A1895601). The initial experiment tested 10, 50 or 500 nM [Pyr1]-Apelin-13 (MedChemExpress, HY-P1033) on days 0 and 1. Untreated cells and cells receiving 6 µM CHIR99021 (Cayman Chemical, 13122) on both days served as comparison conditions. A second experiment tested 2 μM CHIR99021 on days 0 and 1 or on day 0 alone, with or without 50 nM Apelin added on day 1. Medium was replaced on day 1, and Brachyury expression was measured on day 2. Each condition comprised three replicates.

Cells were dissociated with 0.05% trypsin-EDTA (Gibco, 25-300-062), neutralized with serum-containing medium, fixed in 1% paraformaldehyde (Electron Microscopy Sciences, 15710-S) for 20 min at room temperature, and permeabilized in 90% methanol at −20°C overnight. Cells were washed three times with PBS containing 2.5% BSA and 0.1% Triton X-100 (Fisher Scientific, BP151-500), then incubated overnight at 4°C with anti-Brachyury antibody (R&D Systems, AF2085; 1:100). After washing, cells were stained with chicken anti-goat IgG–Alexa Fluor 488 (Invitrogen, A-21467; 1:1,000), washed twice, and resuspended in PBS containing 2.5% BSA. Flow cytometry was performed using a BD Accuri C6 Plus and analyzed using FlowJo 10.8.1. The Brachyury-positive fraction was calculated within the forward- and side-scatter cell gate.

Statistical analyses were performed in GraphPad Prism 11. Data are presented as individual measurements and mean ± SD. An ordinary one-way ANOVA followed by Tukey's multiple-comparison test compared untreated cells with the three Apelin-only conditions. The 6 μM CHIR99021 positive control was displayed separately. An ordinary two-way ANOVA tested the effects of the low-dose CHIR99021 schedule, Apelin treatment and their interaction. The two within-schedule comparisons of Apelin treatment were adjusted together using Šídák's procedure.

**S4. Repeated autonomous searches for tumor antigens**

We initiated three autonomous AI analyses for multiple myeloma (MM) and three for pancreatic ductal adenocarcinoma (PDAC). Each disease-specific prompt asked the agent to select and collect public single-cell data, identify malignant cells, evaluate expression across patients, and prioritize cell-surface proteins for potential targeting by engineered immune cells. The agent was also asked to examine normal-cell expression and assess selected candidates in another dataset. Disease-specific instructions were reused across runs, while dataset selection and implementation were left to the agent.

**S4.1 Multiple myeloma**

MM Runs 1 and 2 used GSE193531 and GSE161801 in opposite discovery and validation roles; Run 3 used GSE232988 for discovery and GSE193531 for validation [2–4]. The discovery populations comprised 10,754 malignant cells from eight donors, 53,729 from 19 donors and

7,228 from 22 donors, respectively. Runs 1 and 2 used the original neoplastic or clonal labels with supporting expression information. Run 3 combined a plasma-cell expression program with a dominant expanded B-cell receptor clone.

Run 1 required at least 500 counts and 200 detected genes per cell, a mitochondrial fraction of 30% or less, and a library-size outlier assessment using four scaled median absolute deviations. Run 2 used the same count and gene minima with a three-median-absolute-deviation lower-outlier rule and an additional joint assessment of mitochondrial fraction. Run 3 used library-specific outlier criteria together with the original low-quality flags. Complete quality-control definitions and retained sample counts are provided in the run-comparison tables.

A gene was considered detected in a cell when its raw count was at least one. In Run 1, a positive donor required at least 30 malignant cells, detection in at least 20% of those cells, and at least one count per million in the summed donor expression profile. Candidate inclusion required positive expression in at least 60% of donors. Run 2 required at least 30 malignant cells per donor, detection in at least 25% of cells, and positive expression in at least 70% of donors. Run 3 required at least 50 malignant cells per donor, at least 25% detection, at least 10 counts per million, positive expression in at least 70% of donors, and at least seven positive donors.

All three runs required evidence supporting cell-surface localization. Run 1 ranked genes in order of donor prevalence, cell detection and expression abundance. Runs 2 and 3 used weighted scores incorporating these quantities. Normal expression was evaluated alongside the rankings; increased expression relative to normal plasma cells was not an initial inclusion requirement. Candidate selection and external expression assessment were completed before comparison with the specified set of established MM antigens.

#### S4.2 Pancreatic ductal adenocarcinoma

The three PDAC analyses searched for public human single-cell datasets with count matrices and sufficient sample information to separate discovery from additional tumor and normal-reference analyses. Run 1 used GSE155698 for discovery, GSE154778 and GSE205013 for additional tumor comparisons, and GSE81547 as an adult normal-pancreas reference. Run 2 used GSE229413 for its primary analysis, including 16 tumor libraries from 15 patients, three adjacent-tissue samples and 11 organ-donor libraries from six donors. GSE229413 contains reprocessed Steele tumor data, so its tumor arm and GSE155698 were treated as one underlying study. GSE212966, GSE205013 and GSE154778 supplied additional quality-control and annotation analyses in Run 2. Run 3 used GSE155698 for discovery and GSE154778 and GSE111672 for additional comparisons. The retained-cell totals were 179,441, 279,924 and 76,221, respectively. The Run 1 total excludes the separately analyzed adult normal reference; the Run 2 total excludes the original-Steele technical comparison; and the Run 3 total includes the selected metastatic samples in its broader cell totals. Dataset roles and run-level cell counts are summarized in Supplementary Table S1 [5–9].

Quality filtering was selected within each run. Run 1 used joint quality flags based on three median absolute deviations, together with a mixture-model assessment of damaged cells. Run 2 used dataset-specific joint mitochondrial and gene-complexity rules. In its primary dataset, a cell was excluded when its mitochondrial percentage exceeded the library median by five unscaled median absolute deviations and its detected-gene count was below the library median. Three- and seven-deviation alternatives were also examined. Run 3 retained cells with at least 200 detected genes for 10x data or 300 for inDrop data and mitochondrial percentages no greater than 30%. Scrublet assessed possible doublets in each run. Runs 1 and 3 excluded predicted doublets, whereas Run 2 retained predictions as flags and considered additional mixed-lineage evidence. The expected doublet rates were adjusted by loading estimates in Run 1, set to 6% in Run 2 and set to 5% in Run 3. No batch integration was applied in these PDAC analyses.

Malignant-cell assessment combined epithelial markers, tumor origin and RNA-derived copy-number patterns, with different operational criteria in each run. Run 1 retained 2,206 eligible malignant cells from seven donors and 674 adjacent ductal cells from three donors. Its confirmatory comparison required five donors per group and at least 30 cells per donor-compartment. Run 3 identified 374 malignant cells in its primary discovery dataset, of which 294 from two donors met the minimum of 20 cells per donor. Its confirmatory analysis required at least three donors per group; three adjacent ductal donors contributed 579 comparison cells.

Run 2 used infercnvpy 0.6.1 with a primary window of 100 genes, a step of ten genes and clipping at three. Eligible libraries required at least 5,000 coordinate-mapped genes and at least 100 genes per autosome. At least two reference lineages with 60 cells each supplied equal training and held-out subsets. Candidate copy-number regions required deviations above the held-out reference 99th percentile, at least 70% directional agreement, a span of at least 10 Mb containing 100 mapped genes, and support on two autosomes. Individual cells also had to satisfy the corresponding regional and burden criteria. Calls were evaluated with windows of 50 and 200 genes, a second reference split, alternative denoising and an immune-only reference. Matching signed regions required at least 50% reciprocal overlap. None of the primary-setting malignant calls remained supported across all applicable checks, so no malignant-versus-normal target model was fitted.

#### S4.3 PDAC expression criteria and candidate comparison

Run 1 assessed genes with membrane annotations using malignant-cell detection, patient prevalence and nonmalignant expression. Its exploratory coverage rule required a median malignant detection fraction of at least 25%, with at least 60% of donors having expression in 20% or more of their malignant cells. The low-background rule required a one-sided 95% bootstrap upper bound of no more than 5% in the evaluated nonmalignant populations. The 20 reported surface-associated genes were retained for examination of why they failed the combined criteria. Run 3 required median malignant detection of at least 50%, expression in at least 30% of malignant cells in at least 70% of donors, and a descriptive malignant-versus-ductal expression ratio threshold before retaining 20 exploratory genes. Its selectivity assessment required every eligible nonmalignant donor fraction to be at most 10% and a median malignant-minus-nonmalignant difference of at least 40 percentage points. All 20 genes exceeded the nonmalignant-expression limit. Six retained expression coverage in both additional primary-tumor comparisons: GPRC5A, CEACAM6, LSR, F11R, DSG2 and CLDN4.

The intended Run 2 target comparison required five malignant donors with at least 50 cells each. Because malignant-cell eligibility failed, no enrichment, patient-prevalence or candidate-validation test was performed. Descriptive reference aggregation nevertheless included 57,687 cells in 94 donor-compartment groups. Raw counts and gene-detection numerators were summed by donor and compartment, combining technical libraries. Ductal summaries required at least 50 cells per donor and other compartments required at least 30. Organ-donor, adjacent-tissue and tumor-associated reference compartments were kept separate. Six organ donors met the cell-number minimum for ductal, acinar, endocrine, fibroblast and endothelial summaries; three met it for immune cells. Pericyte identity remained unresolved. These summaries describe the available normal-reference expression data and do not contribute a PDAC candidate ranking.

Candidate membership was compared using the final reported surface lists and recorded Ensembl identifiers. Runs 1 and 3 contributed 20 genes each and had no shared gene at top 5, 10 or 20. The corresponding Jaccard values were zero. Run 2 did not produce a candidate list, so pairwise comparisons involving Run 2 and three-way overlap were unavailable. The combined primary and secondary lists contained 70 distinct genes: 30 from Run 1 and 40 from

Run 3. Secondary lists were kept separate from the surface-antigen comparison because their localization and prioritization criteria differed.

#### S4.4 Comparison across runs

A separately prompted AI analysis compared candidate membership and ranks using the lists generated by each run. For each disease, recurrence was the number of runs in which a gene appeared. Pairwise overlap among the top-ranked genes was summarized using the number shared and the Jaccard index, calculated as the number of shared genes divided by the total number of distinct genes in the compared lists. Gene ranks were taken from the candidate lists, with genes absent from a list left unranked. MM recovery of established targets used four antigens as its denominator. These analyses summarize agreement among the computational runs.

### S5. Blinded and context-informed reanalysis of lung single-cell data

#### S5.1 Dataset preparation and analysis conditions

We used the IPF and nonfibrotic-control subset of GSE135893 [10], comprising 89,326 cells from 22 donors and 31 sequencing libraries. This subset included 31,644 cells from ten controls in 12 libraries and 57,682 cells from 12 IPF donors in 19 libraries. The full source study included additional pulmonary-fibrosis diagnoses and 114,396 cells from 30 donors.

Before the blind analysis, disease groups were renamed Group A and Group B. Donor, library, specimen-source, processing-site, flowcell and chemistry identifiers were replaced with anonymous codes, and accessions, original sample names, cell annotations and identifying study information were withheld. We initiated the blind analysis in a separate account, with personalization and memory disabled and Work reset. The prompt prohibited retrieval of prior conversations or analyses, identification of the source study, and disease-specific literature searches before the analysis was complete. General biological knowledge and software documentation could be used. The context-informed analysis received the same anonymized data with the human lung, IPF and control labels, but without the publication, original annotations or blind-analysis results. Each analysis was finalized before comparison with the source publication.

Both prompts requested quality control, cell annotation, donor-level comparisons, analysis of gene programs, defined as groups of genes involved in a related biological process, and investigation of relationships between cell populations. Processing site and specimen source were represented as separate variables, with two processing sites and three specimen-source categories. Both conditions occurred at each processing site. One specimen source included seven controls and seven IPF donors, whereas the other two source categories were condition specific. Source-restricted analyses were therefore used to assess whether findings depended on this sampling structure.

#### S5.2 Computational analysis

Both analyses required at least 1,000 detected genes and less than 20% mitochondrial RNA per cell. The context-informed analysis additionally removed extreme library-specific count outliers. The blind analysis retained 87,705 cells and used log-normalized counts per 10,000, 3,500 donor-aware variable genes, 50 principal components, and a 20-neighbor graph based on the first 40 components. The context-informed analysis retained 87,698 cells and used 2,500 library-aware variable genes, 40 principal components and 20 neighbors. Both used Leiden clustering, UMAP and marker-based annotation.

For gene-expression comparisons, raw counts were summed within each cell population and donor to obtain donor-level expression profiles, also termed pseudobulk profiles. Both analyses used ordinary least-squares models with disease-group and processing-site terms and heteroscedasticity-consistent HC3 standard errors. The blind analysis used log2(CPM + 0.5), requiring at least 20 cells in each of at least five donors per group. The context-informed analysis required at least four donors per group and used median-ratio normalization, with total-count normalization when a size factor could not be estimated. Genes were eligible when they reached at least one count per million in four donors.

The Benjamini–Hochberg procedure controlled the false discovery rate across 128,323 cell-population–gene tests in the blind analysis and 230,062 gene–stratum tests in the context-informed analysis. The latter included overlapping cell identities and lineages. Cell-composition comparisons used donor-level cell fractions in the blind analysis and empirical-logit-transformed fractions in the context-informed analysis. Gene programs, cell states and associations between cell populations were evaluated in separate multiple-testing families. Additional analyses examined the effects of specimen source, stricter cell filtering, annotation choices, normalization and individual donors.

#### S5.3 Comparison with the source publication

After completion of the two analyses, a separately prompted AI comparison examined correspondence with 42 findings from the original paper and its supplementary results. A finding was directly comparable when the required cell populations, conditions and measurements were represented in the reanalyzed subset, partially comparable when only part of that information was available, and unavailable when the result depended on excluded diagnoses or another assay. A separate comparison asked whether each relationship was discussed in the main text, discussed in the supplement, represented by gene-level tables without interpretation of the same relationship, absent as a specific relationship, or unresolved. The complete correspondence tables provide the supporting excerpts and category definitions.

**S5.4 Tests in another patient dataset**

A separately prompted AI analysis independently selected 13 additional questions from the blind and context-informed findings and evaluated them using GSE227136 [11]. Matching against all 30 donor identifiers in the full discovery study excluded previously analyzed donors, leaving 70 donors, comprising 28 IPF and 42 controls, in 91 libraries. The source studies shared a collection network, so the external analysis tested different patient records within that network. Two myeloid-state comparisons and three additional technical controls were evaluated separately from the 13-question set.

Cells were retained with 500–50,000 counts, 200–6,000 detected genes and at most 20% mitochondrial RNA. The AI agent independently established the cell eligibility requirements, gene sets, predicted directions and statistical testing criteria before examining the corresponding expression outcomes. Cell identities were assigned using predefined marker panels, and each question had a minimum cell-coverage requirement per donor. Gene-program analyses used log2(CPM + 1) donor-level expression, standardized across donors before combining the specified genes into a score. This places genes on a comparable scale so that the score reflects their combined expression pattern. The analysis tested whether donors with higher gene-program scores in one cell population also had higher scores in another, after adjustment for diagnosis, a site variable derived from donor identifiers, mean log cell counts, mean mitochondrial fraction and library number. The predefined direction was tested using one-sided tests, and approximate two-sided Fisher confidence intervals were reported for the correlation coefficients.

CRYM expression in fibroblasts was assessed using a negative-binomial donor-count model accounting for the total fibroblast RNA counts and collection site. The total-count offset compares CRYM abundance relative to the amount of fibroblast RNA measured for each donor. A complementary model tested the proportion of CRYM-positive fibroblasts using empirical-logit transformation and equal-donor ordinary least squares with HC3 standard errors. Both changes had to support the predicted direction; the larger of the two component P values was used for multiple-testing adjustment. Benjamini–Hochberg correction was applied to six prespecified positions in each of three testing families. Positions corresponding to untestable questions or cell-identity assessment were assigned P = 1, preserving the original family sizes.

A question was supported in the initial test when it met the predefined direction and statistical threshold. The AI analysis then assessed whether the result remained supported when cell filtering, cell definitions or individual-donor inclusion changed. Results that established a rare cell population without sufficient group coverage were reported as detection of that population rather than a disease-associated change. We also retained a separate classification for signals whose biological interpretation could not be resolved from technical effects. The selected questions and overall outcomes are summarized in Supplementary Table S6. Detailed analysis records, donor-level measurements and complete external-test results are available in the computational repository (Data and code availability).

**S6. Integration of public pancreatic cancer data**

In this autonomous investigation, the AI agent selected and integrated public transcriptomic, protein, functional and normal-tissue data to evaluate pancreatic cancer targets, including intracellular as well as cell-surface candidates. RNA discovery used processed microarray data from 60 paired tumor and adjacent-tissue specimens in GSE62452 [12]. Probes were mapped unambiguously to genes, and paired t tests were applied to the log-scale expression values. Eligibility required a mean tumor–adjacent log2 difference of at least 1 and Benjamini–Hochberg $q \le 0.05$. External expression testing used 118 tumors and 13 controls from GSE62165, with Welch's tests and thresholds of log2 difference $\ge 0.5$ and $q \le 0.05$ [13]. The 45 GSE62452 pairs also present in GSE28735 and the remaining 15 pairs were examined as subsets of the same discovery study.

Single-nucleus RNA data from 108,964 nuclei from 18 untreated GSE202051 tumor donors were used to examine expression across cell compartments [14]. Counts were summed within each donor and compartment, retaining the original annotations and requiring at least 20 nuclei per compartment and five paired donors per contrast. Paired bulk-protein analysis used up to 62 CPTAC donors, with the number of complete pairs determined separately for each gene [15]. Missing protein values were not imputed. Strong protein support required a mean paired difference of at least 0.5 on the processed log2 scale, $q \le 0.05$ and at least ten complete pairs. Methylation, copy-number, variant and protein-adjusted phosphosite analyses provided additional molecular context from overlapping CPTAC patients.

Cancer-cell dependency, meaning reduced growth or survival after gene disruption, was assessed using the BAGEL2 calls provided with the source CRISPR screens [16,17]. Model and donor reconciliation yielded 20 donor-level PDAC model groups and 849 groups from other cancers. Selective dependency required data from at least ten PDAC model groups, a dependency-prevalence difference of at least 0.20, and a one-sided Fisher-test $q \leq 0.05$ after adjustment across 17,645 genes. Four pancreatic-adenocarcinoma organoid models supplied additional functional observations. Expression in normal tissues was evaluated using RNA and protein references, while Human Protein Atlas and UniProt annotations informed cellular localization and candidate properties. Normal expression was classified as high when a specified critical tissue had at least 100 normalized transcripts per million or medium/high protein staining. Detectable expression below this category was defined as at least 10 normalized transcripts per million. Critical tissues included heart, brain, lung, liver, kidney, bone marrow, esophagus, stomach, small intestine, colon and pancreas.

The combined candidate set comprised genes meeting the RNA or functional criteria. Each candidate was assessed across ten evidence domains: PDAC association, malignant-cell specificity, patient prevalence, agreement across datasets, protein abundance, spatial support, functional dependency, experimental perturbation, genomic or regulatory support, and biological plausibility. Measured evidence received 0–2 points according to the domain-specific definitions; missing measurements retained an explicit missing value and received no positive score. Supplementary Table S8 summarizes the evidence categories and scoring definitions. The complete component scores and implemented scoring rules are provided in the deposited evidence matrix and analysis code. Ranking prioritized the level of supporting evidence, the summed biological-support score, the number of independent patient datasets providing evidence and major contradictory findings. Shared scores received shared ranks. Normal-tissue expression and properties relevant to intervention were evaluated separately from positive molecular support. Candidates were classified according to the amount of evidence available for further therapeutic testing. The higher categories required evidence for the target protein in the relevant cell compartment and an intervention range that could affect cancer cells while sparing normal cells.

To examine how different measurements affected prioritization, we compared rankings obtained as evidence types were added while holding the 213-candidate set constant. The comparison began with RNA support and successively included single-nucleus, protein, functional, normal-reference and remaining evidence. This assessed the contribution of the included data to the final ranking. Target-specific therapeutic literature review was performed after candidate selection and covered the first 20 displayed candidates and the six candidates identified through selective dependency. A separate comparison examined overlap with the specified set of established treatment mechanisms.

**S7. Pancreatic spatial-transcriptomic viewer**

The viewer used a publicly available 10x Genomics Xenium pancreatic adenocarcinoma section containing 190,965 segmented cells [18]. Cell centroids were displayed at their measured x and y coordinates in micrometers. Six display groups combined the 16 provider cell annotations, with three unassigned cells retained separately. Cell-group filters and gene overlays enabled interactive inspection of their tissue distribution.

The ten available overlays displayed measured RNA counts for EPCAM, CFTR, ANPEP, PPY, CD163, CD3E, MS4A1, CTHRC1, VWF and CXCL9. The assay used a targeted panel of 377 predesigned and 97 additional genes. Dataset attribution and the CC BY 4.0 license accompany the viewer and its derivative image. The interactive viewer is available at https://scintillating-jin-8e1ba7.netlify.app/model-02.

### Data and code availability

Computational data, code, available prompts and analysis records from Parts 1–4 and the spatial resource are deposited in Zenodo at https://doi.org/10.5281/zenodo.23290610. The record includes a README and file index describing the deposited computational materials.

### Supplementary Prompt Summary

We began with a biological aim for each analysis, used AI assistance to develop the detailed instructions, and refined the resulting prompts before submission. The table below summarizes the purpose of each prompt group. Available prompt records and associated documentation are provided in the computational repository at https://doi.org/10.5281/zenodo.23290610.

| **Prompt group** | **Purpose** |
|---|---|
| P1. Interactive hPSC analysis | Identify signals associated with early mesendoderm differentiation and guide data analysis step by step. |

| Prompt group | Purpose |
| --- | --- |
| P2. Autonomous PDAC analysis | Collect and analyze public single-cell data to evaluate tumor-associated surface antigens across patients. |
| P3. Autonomous MM analysis | Evaluate candidate surface antigens, their expression in another dataset and recovery of established treatment antigens. |
| P4. Repeated-run comparisons | Compare the datasets, analytical choices, candidate lists, rankings and biological conclusions from three runs per disease. |
| P5. Blind analysis | Analyze anonymized expression data without the source publication, accession, tissue or diagnosis labels. |
| P6. Context-informed analysis | Analyze the same data with tissue and disease labels while withholding the publication and blind-analysis results. |
| P7. Publication comparison | Compare the completed analyses with the original paper and identify additional relationships for further study. |
| P8. External testing | Test the selected relationships using other patient records and specified statistical criteria. |
| P9. Broader PDAC investigation | Integrate available molecular, functional and normal-tissue evidence for candidate prioritization. |
| P10. Spatial viewer | Create an interactive view of pancreatic cell locations and selected measured genes. |

For the lung analyses, a separate provenance table distinguishes archived run-associated instructions from later supplied copies. The archived blind-analysis instructions include an additional context-isolation preamble that is absent from the later copy. Both records are retained in the repository.

### Supplementary Table Legends

**Supplementary Table S1. Datasets and analysis populations.** Public data accessions, source publications, measurement types, analyzed conditions, donor or specimen counts, cell counts and the role of each dataset in candidate discovery, testing in another dataset or biological interpretation. Counts refer to the population used for each analysis and may describe a subset of the source study.

**Supplementary Table S2. Computational settings for repeated autonomous analyses.** Dataset selection, cell-quality filters, malignant-cell definitions, donor requirements, expression thresholds, ranking methods and testing criteria for each MM and PDAC run.

**Supplementary Table S3. Candidate membership and recurrence across autonomous runs.** Complete candidate lists, ranks within each run, the number of runs selecting each gene, and overlap between candidate lists. MM and PDAC are reported separately, with the compared list types specified.

**Supplementary Table S4. Recovery of established MM antigens.** Selection, candidate rank and RNA-expression support in another dataset for the four reference antigens in each run. BCMA is represented by TNFRSF17. An absent rank means that the run did not select that antigen.

**Supplementary Table S5. Comparison of lung analyses with the source publication.** The 42 source findings, whether they could be examined in the reanalyzed data subset, and the corresponding findings from each analysis. Additional entries distinguish relationships discussed in the publication from gene-level results available in its tables and relationships developed through the reanalysis.

**Supplementary Table S6. Additional lung relationships and tests in other donors.** Summary of the 13 selected questions, relevant cell populations, numbers of eligible donors, effect estimates, confidence intervals, adjusted P values and overall outcomes. Outcomes distinguish supported associations, detection of the predicted cell population, possible technical signals and insufficient data for testing. Gene lists and detailed statistical results are available in the deposited Part 3 external-validation records.

**Supplementary Table S7. Donor-level measurements for selected lung relationships.** Gene-expression summaries for the relationship between AT2 and secretory-cell redox programs, and fibroblast CRYM RNA counts and detection percentages. The table also includes diagnostic groups, cell counts and measurements for the selected analyses.

**Supplementary Table S8. Integrated evidence for PDAC candidates.** All 213 candidates, the RNA or functional evidence that qualified them for inclusion, additional molecular support, normal-tissue expression and final rank. The scoring definitions explain the 0–2 points assigned within each evidence category. The table distinguishes the 207 candidates selected through increased tumor RNA from the six selected through preferential effects of gene disruption on PDAC cells. Missing measurements are indicated explicitly.

**Supplementary Table S9. Contribution of evidence types to PDAC prioritization.** Candidate rankings after RNA, cell-type, protein, functional and other evidence were added in sequence, and comparison with the complete ranking. Every stage uses the same 213 genes. The accompanying treatment-mechanism comparison records the reference set and its overlap with the candidates.

**Supplementary Table S10. Related AI approaches for biological data analysis.** Published descriptions of analysis agents, broader biomedical systems, interpretation tools, representation models and evaluation frameworks. Entries summarize tasks, data inputs, interfaces, computing requirements, human involvement, result verification and reported experimental testing.

### Supplementary Figures

A

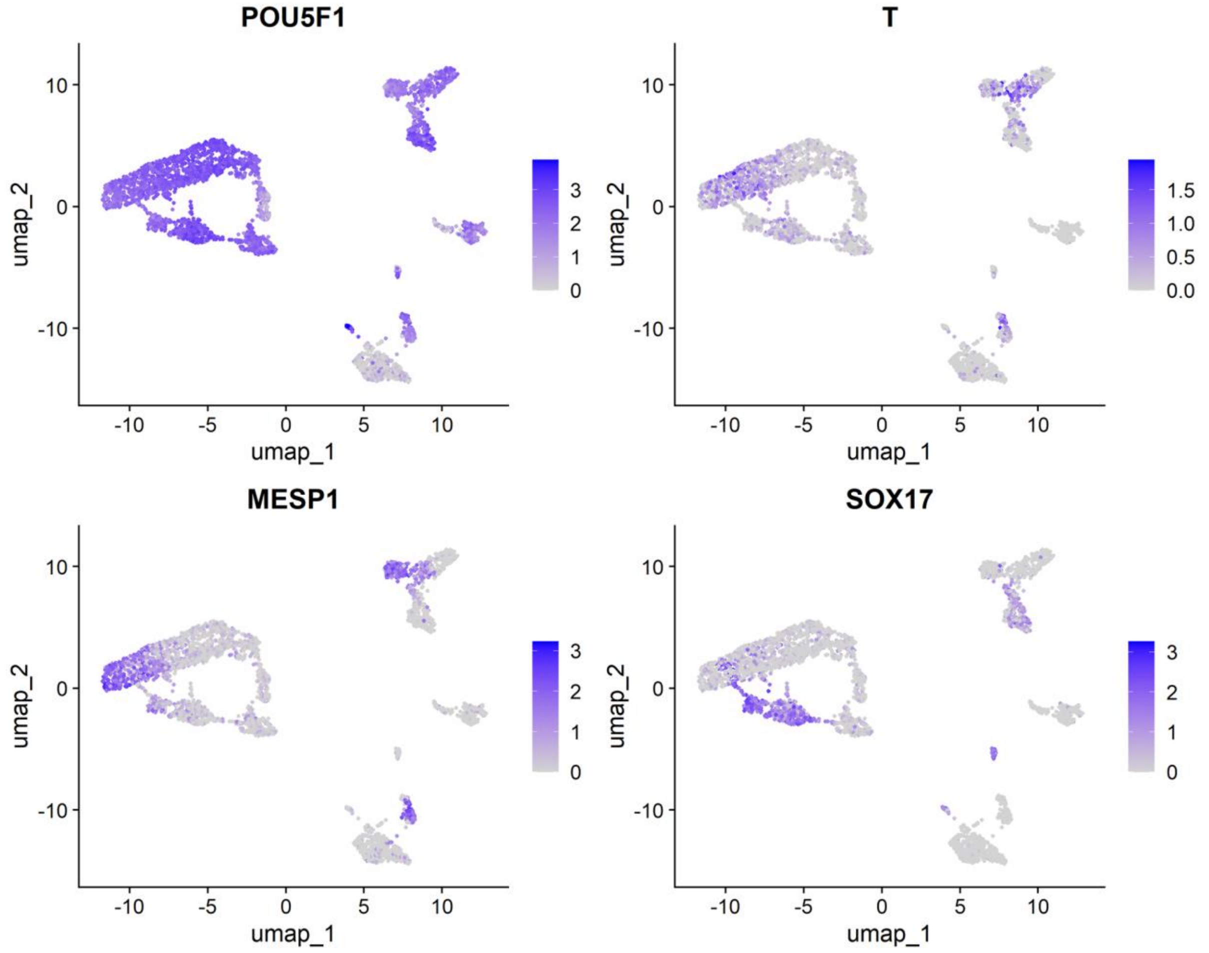


B

| Cell state | Clusters | Cells |
|---|---|---|
| Putative pluripotent | 0, 6, 7 | 1,023 |
| Primitive-streak-like | 1 | 562 |
| Mesoderm | 3, 9 | 496 |

**Supplementary Figure S1. Marker expression and cell populations used for the gastruloid analysis.** (A) Expression of POU5F1, T, MESP1 and SOX17 on the cell map shown in Figure 2A. Each point represents a cell, and cells with similar overall RNA-expression patterns are positioned near one another by UMAP. Purple intensity indicates log-normalized expression of the specified gene. T is the name used for TBXT in the source expression matrix. (B) Clusters and cell counts for the putative pluripotent, primitive-streak-like and mesoderm populations selected for the candidate analysis.

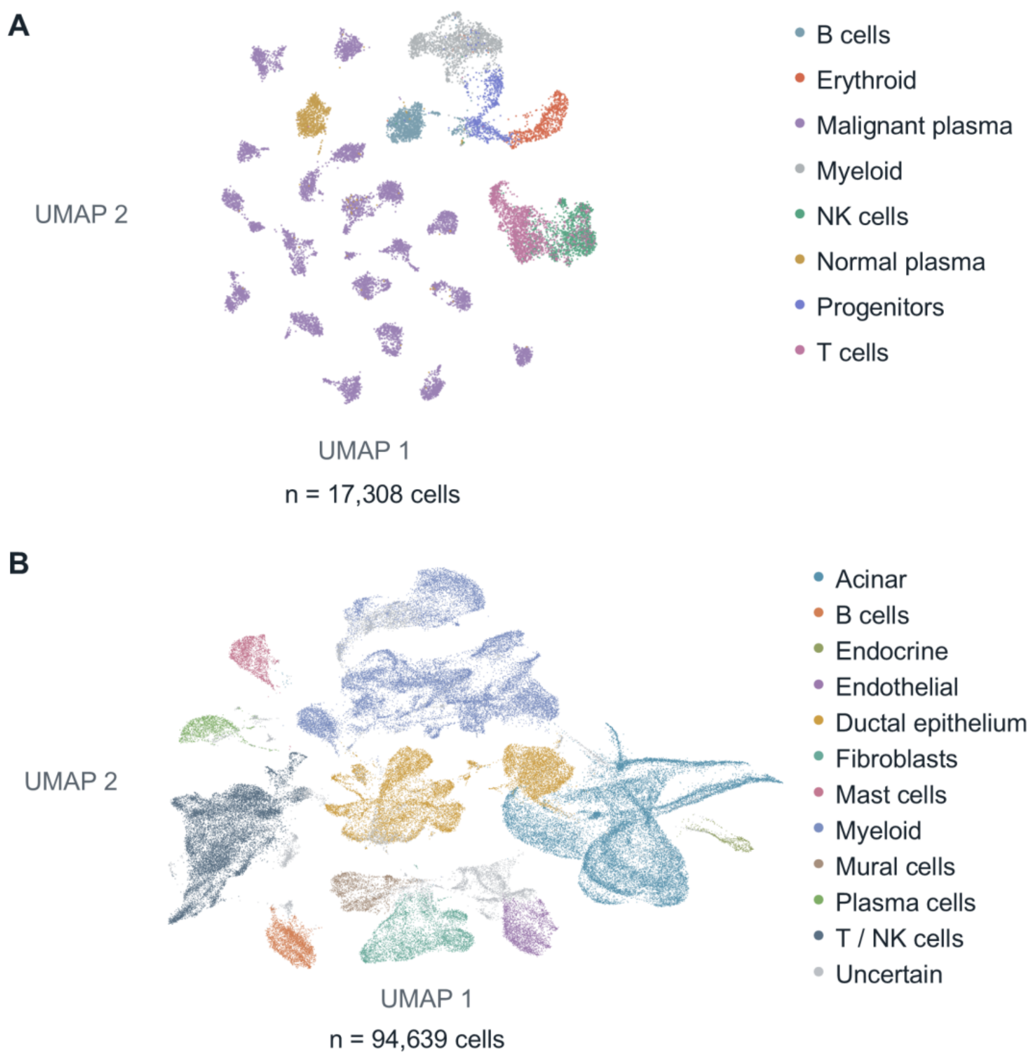


**Supplementary Figure S2. Cell populations identified during autonomous MM and PDAC analysis.** Each point represents a cell, positioned by UMAP according to its RNA-expression pattern and colored by its assigned cell population. (A) MM Run 2, using GSE161801. The displayed subset contains 17,308 cells; candidate analysis used the full eligible population of 53,729 malignant plasma cells from 19 donors. (B) PDAC Run 2, showing all 94,639 retained cells from the primary dataset, GSE229413. This dataset includes tumor, adjacent-pancreas and organ-donor samples. Mural cells include cells with pericyte or smooth-muscle characteristics. MM, multiple myeloma; PDAC, pancreatic ductal adenocarcinoma; NK, natural killer.

A

| Analysis / question | n | External result |
|---|---|---|
| Blind: AT2–endothelial immunoglobulin RNA | 48 | Background RNA recurs |
| Blind: monocyte–lymphoid heat-shock RNA | 45 | Stress RNA recurs |
| Blind: myeloid interferon-response RNA | 0 | No eligible control donors |
| Blind: AT2–secretory redox-related RNA | 47 | $r = 0.830$, $q < 0.001$ |
| Blind: endothelial–macrophage fatty-acid oxidation | 49 | $r = 0.283$, $q = 0.038$* |
| Blind: endothelial–macrophage erythroid RNA | 49 | Background RNA recurs |
| Blind: rare neuroendocrine-like epithelial cells | 70 | 78 coherent cells in 5 donors |
| Context: fibroblast CRYM RNA | 31 | Rate ratio 4.52, $q < 0.001$ |
| Context: endothelial glycolysis-related RNA | 50 | Score +0.445, $q = 0.006$ |
| Context: AT2/macrophage damage-response RNA | 59 | Score +1.553, $q < 0.001$ |
| Context: lymphoid MHC-II-related RNA | 51 | Score +1.018, $q < 0.001$ |
| Context: epithelial–myeloid redox-related RNA | 63 | $r = 0.388$, $q = 0.002$ |
| Context: myeloid–vascular respiratory RNA | 51 | $r = 0.563$, $q < 0.001$ |

B

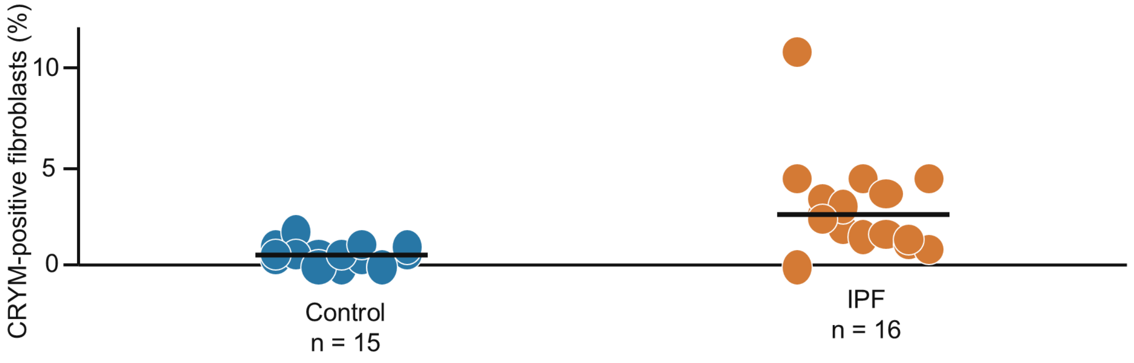


**Supplementary Figure S3. Results for all 13 additional lung questions and CRYM detection in fibroblasts.** (A) Questions generated by the blind or context-informed analysis and tested in a second lung dataset containing 70 donors. Each row summarizes one question; n is the number of donors with sufficient relevant cells for that test. Positive correlation coefficients (r) indicate that donors with higher expression of the specified gene set in one cell population also tended to have higher expression in the other, after adjustment for diagnosis and technical variables. Gene-set scores combine standardized expression of the listed genes; positive score differences indicate higher values in IPF than controls, except for the AT2/macrophage comparison, which tests the difference between the disease-associated changes in these two cell populations. The CRYM rate ratio compares fibroblast CRYM RNA abundance in IPF and control donors after accounting for the total RNA measured. q denotes the P value adjusted for multiple testing. The rare-cell search examined all 70 donors and detected 78 neuroendocrine-like cells in five donors. Stress or background RNA denotes recurring signals that could reflect sample processing or RNA originating outside the assigned cell population. The interferon question could not be tested because no control donors had enough of the required dendritic cells. Summary results for the 13 questions are provided in Supplementary Table S6. *For the fatty-acid-oxidation relationship, the one-sided test supported the predicted positive direction, but the two-sided 95% confidence interval included zero and the result did not remain significant with stricter cell filtering. (B) Percentage of fibroblasts with detected CRYM RNA in 15 control and 16 IPF donors. Each point represents one donor, and horizontal lines show medians. AT2, alveolar type 2; IPF, idiopathic pulmonary fibrosis; MHC-II, major histocompatibility complex class II.

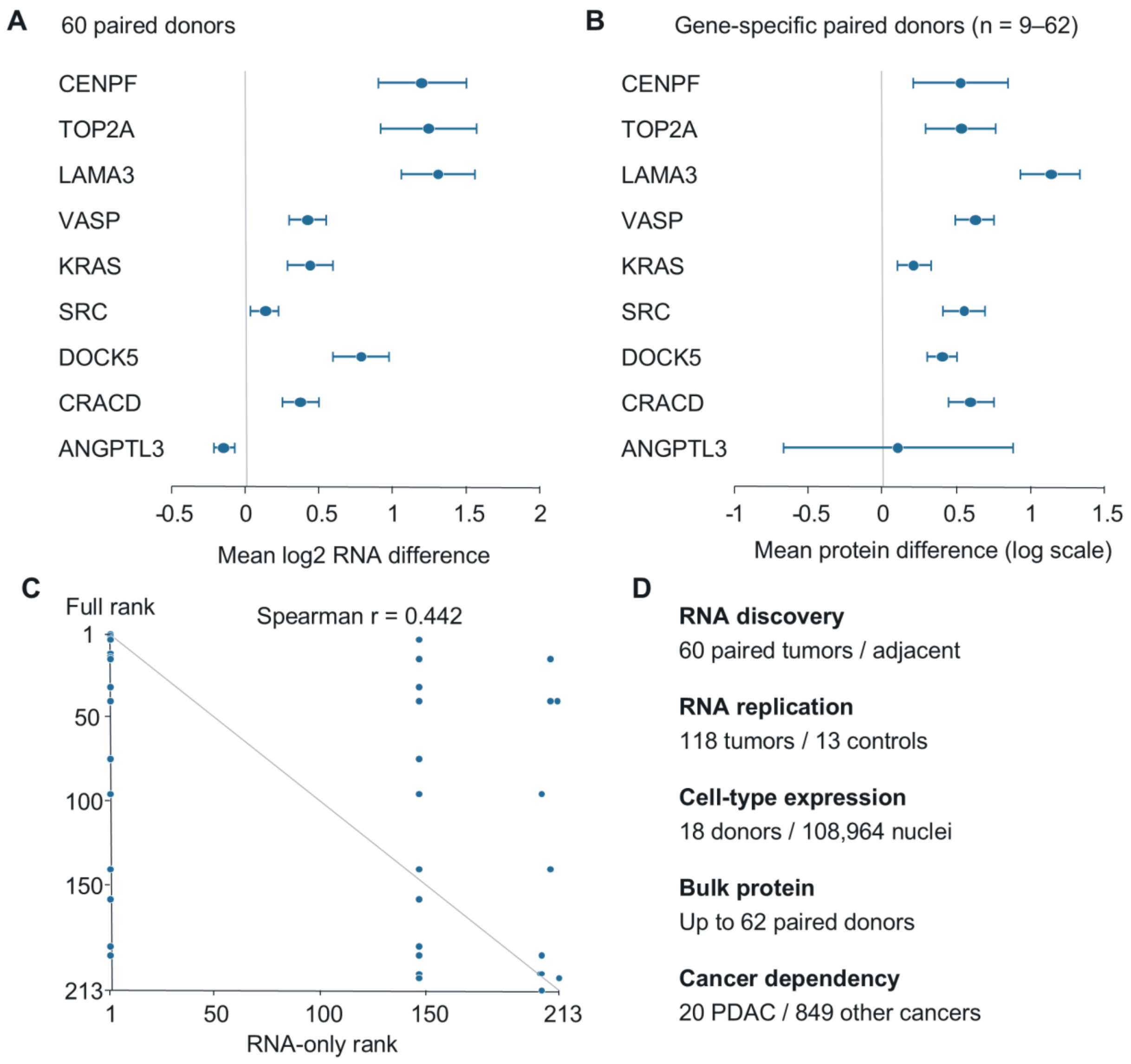


**Supplementary Figure S4. RNA and protein differences and their contribution to pancreatic cancer candidate rankings.** (A) Mean tumor-minus-adjacent-tissue differences in log2 RNA expression for the nine candidates shown in Figure 7B, measured in 60 paired donors from GSE62452. A value of 1 corresponds to an approximately twofold difference in RNA expression. (B) Mean paired tumor-minus-adjacent-tissue protein differences in CPTAC. Points in (A,B) indicate means, and error bars show 95% confidence intervals; values above zero indicate greater abundance in tumors. Complete protein pairs were available for 43 donors for CENPF, 52 for TOP2A, 62 for LAMA3, VASP, KRAS, SRC, DOCK5 and CRACD, and nine for ANGPTL3. (C) Candidate priority using RNA evidence alone versus all included evidence, within the same set of 213 genes. Smaller rank numbers indicate higher priority. Points above the diagonal received higher priority when the additional evidence was included; points below it received lower priority. Genes with equal scores share ranks, so several genes can occupy the same point. Spearman r summarizes agreement between the two rankings. (D) Principal datasets contributing RNA, cell-type, protein and gene-dependency evidence. Dependency describes reduced cancer-cell growth or survival after gene disruption. PDAC, pancreatic ductal adenocarcinoma; CPTAC, Clinical Proteomic Tumor Analysis Consortium.

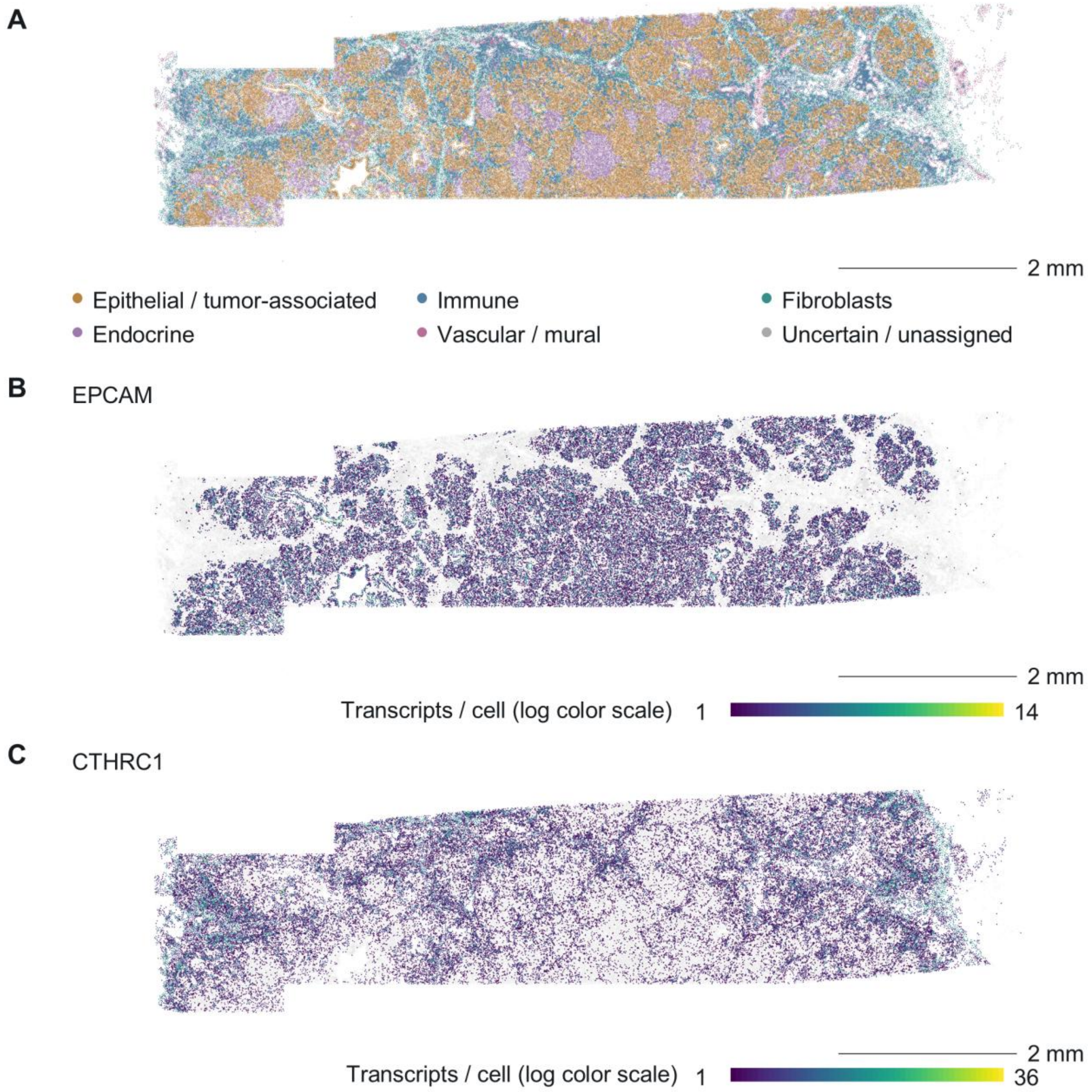

**Supplementary Figure S5. Spatial visualization of a public pancreatic adenocarcinoma dataset.** (A) Spatial distribution of 190,965 segmented cells grouped into six broad cell categories based on the original annotations. (B, C) Spatial RNA expression of EPCAM (B) and CTHRC1 (C) across the same tissue section. Each point represents a segmented cell at its measured tissue coordinates, with colors indicating cell categories in A or assigned transcript counts per cell in B and C. RNA expression is displayed using a logarithmic color scale, with separate ranges for each gene. Scale bars, 2 mm. Data were obtained from the 10x Genomics Xenium pancreatic adenocarcinoma dataset [18].

9. Lin W, Noel P, Borazanci EH, Lee J, Amini A, Han IW, Heo JS, Jameson GS, Fraser C, Steinbach M, Woo Y, Fong Y, Cridebring D, Von Hoff DD, Park JO, Han H. Single-cell transcriptome analysis

of tumor and stromal compartments of pancreatic ductal adenocarcinoma primary tumors and metastatic lesions. Genome Med. 2020;12(1):80. doi:10.1186/s13073-020-00776-9

10. Habermann AC, Gutierrez AJ, Bui LT, Yahn SL, Winters NI, Calvi CL, Peter L, Chung MI, Taylor CJ, Jetter C, Raju L, Roberson J, Ding G, Wood L, Sucre JMS, Richmond BW, Serezani AP, McDonnell WJ, Mallal SB, Bacchetta MJ, Loyd JE, Shaver CM, Ware LB, Bremner R, Walia R, Blackwell TS, Banovich NE, Kropski JA. Single-cell RNA sequencing reveals profibrotic roles of distinct epithelial and mesenchymal lineages in pulmonary fibrosis. Sci Adv. 2020;6(28):eaba1972. doi:10.1126/sciadv.aba1972

11. Natri HM, Del Azodi CB, Peter L, Taylor CJ, Chugh S, Kendle R, Chung M i, Flaherty DK, Matlock BK, Calvi CL, Blackwell TS, Ware LB, Bacchetta M, Walia R, Shaver CM, Kropski JA, McCarthy DJ, Banovich NE. Cell-type-specific and disease-associated expression quantitative trait loci in the human lung. Nat Genet. 2024;56(4):595–604. doi:10.1038/s41588-024-01702-0

12. Yang S, He P, Wang J, Schetter A, Tang W, Funamizu N, Yanaga K, Uwagawa T, Satoskar AR, Gaedcke J, Bernhardt M, Ghadimi BM, Gaida MM, Bergmann F, Werner J, Ried T, Hanna N, Alexander HR, Hussain SP. A Novel MIF Signaling Pathway Drives the Malignant Character of Pancreatic Cancer by Targeting NR3C2. Cancer Res. 2016;76(13):3838–50. doi:10.1158/0008-5472.can-15-2841

13. Janky R, Binda MM, Allemeersch J, Van den broeck A, Govaere O, Swinnen J V, Roskams T, Aerts S, Topal B. Prognostic relevance of molecular subtypes and master regulators in pancreatic ductal adenocarcinoma. BMC Cancer. 2016;16(1):632. doi:10.1186/s12885-016-2540-6

14. Hwang WL, Jagadeesh KA, Guo JA, Hoffman HI, Yadollahpour P, Reeves JW, Mohan R, Drokhlyansky E, Van Wittenberghe N, Ashenberg O, Farhi SL, Schapiro D, Divakar P, Miller E, Zollinger DR, Eng G, Schenkel JM, Su J, Shiau C, Yu P, Freed-Pastor WA, Abbondanza D, Mehta A, Gould J, Lambden C, Porter CBM, Tsankov A, Dionne D, Waldman J, Cuoco MS, Nguyen L, Delorey T, Phillips D, Barth JL, Kem M, Rodrigues C, Ciprani D, Roldan J, Zelga P, Jorgji V, Chen JH, Ely Z, Zhao D, Fuhrman K, Fropf R, Beechem JM, Loeffler JS, Ryan DP, Weekes CD, Ferrone CR, Qadan M, Aryee MJ, Jain RK, Neuberg DS, Wo JY, Hong TS, Xavier R, Aguirre AJ, Rozenblatt-Rosen O, Mino-Kenudson M, Castillo CF del, Liss AS, Ting DT, Jacks T, Regev A. Single-nucleus and spatial transcriptome profiling of pancreatic cancer identifies multicellular dynamics associated with neoadjuvant treatment. Nat Genet. 2022;54(8):1178–91. doi:10.1038/s41588-022-01134-8

15. Cao L, Huang C, Cui Zhou D, Hu Y, Lih TM, Savage SR, Krug K, Clark DJ, Schnaubelt M, Chen L, da Veiga Leprevost F, Eguez RV, Yang W, Pan J, Wen B, Dou Y, Jiang W, Liao Y, Shi Z, Terekhanova N V, Cao S, Lu RJH, Li Y, Liu R, Zhu H, Ronning P, Wu Y, Wyczalkowski MA, Easwaran H, Danilova L, Mer AS, Yoo S, Wang JM, Liu W, Haibe-Kains B, Thiagarajan M, Jewell SD, Hostetter G, Newton CJ, Li QK, Roehrl MH, Fenyö D, Wang P, Nesvizhskii AI, Mani DR, Omenn GS, Boja ES, Mesri M, Robles AI, Rodriguez H, Bathe OF, Chan DW, Hruban RH, Ding L, Zhang B, Zhang H, Amin M, An E, Ayad C, Bauer T, Birger C, Birrer MJ, Boca SM, Bocik W, Borucki M, Cai S, Carr SA, Cerda S, Chen H, Chen S, Chesla D, Chinnaiyan AM, Colaprico A, Cottingham S, Derejska M, Dhanasekaran SM, Domagalski MJ, Druker BJ, Duffy E, Dyer MA, Edwards NJ, Ellis MJ, Eschbacher J, Francis A, Francis J, Gabriel S, Gabrovski N, Gardner J, Getz G, Gillette MA, Goldthwaite CA, Grady P, Guo S, Hariharan P, Hiltke T, Hindenach B, Hoadley KA, Huang J, Jones CD, Ketchum KA, Kinsinger CR, Koziak JM, Kusnierz K, Liu T, Long J, Mallery D, Mareedu S, Matteotti R, Maunganidze N, McGarvey PB, Minoo P, Paklina O V, Paulovich AG, Payne SH, Potapova O, Pruetz B, Qi L, Roche N, Rodland KD, Rohrer DC, Schadt

EE, Shabunin A V, Shelton T, Shutack Y, Singh S, Smith M, Smith RD, Sokoll LJ, Suh J, Thangudu RR, Tsang SX, Um KS, Valley DR, Vatanian N, Wang W, Wilson GD, Wiznerowicz M, Zhang Z, Zhao G. Proteogenomic characterization of pancreatic ductal adenocarcinoma. Cell. 2021;184(19):5031-5052.e26. doi:10.1016/j.cell.2021.08.023

16. Pacini C, Duncan E, Gonçalves E, Gilbert J, Bhosle S, Horswell S, Karakoc E, Lightfoot H, Curry E, Muyas F, Bouaboula M, Pedamallu CS, Cortes-Ciriano I, Behan FM, Zalmas LP, Barthorpe A, Francies H, Rowley S, Pollard J, Beltrao P, Parts L, Iorio F, Garnett MJ. A comprehensive clinically informed map of dependencies in cancer cells and framework for target prioritization. Cancer Cell. 2024;42(2):301-316.e9. doi:10.1016/j.ccell.2023.12.016

17. Kim E, Hart T. Improved analysis of CRISPR fitness screens and reduced off-target effects with the BAGEL2 gene essentiality classifier. Genome Med. 2021;13(1):2. doi:10.1186/s13073-020-00809-3

18. 10x Genomics. Pancreatic Cancer with Xenium Human Multi-Tissue and Cancer Panel [Internet]. 10x Genomics; 2023. Available from: https://www.10xgenomics.com/datasets/pancreatic-cancer-with-xenium-human-multi-tissue-and-cancer-panel-1-standard
