## Supplementary Tables for "General Purpose AI for Public Data Analysis and Experimental Hypothesis Generation": Supplementary_Table_S4.pdf

### Supplementary Table S4. Recovery of established MM antigens

Selection, candidate rank and RNA-expression support in another dataset for four reference antigens across three independent analyses. BCMA is encoded by TNFRSF17.

#### A. Antigen selection and expression support

| Antigen / gene | Run 1 | Run 2 | Run 3 | Selected in |
|---|---|---|---|---|
| BCMA<br>TNFRSF17 | **Rank 11**<br>RNA support | **Rank 8**<br>RNA support | **Rank 12**<br>RNA support | 3/3 runs |
| CD38 | **Rank 19**<br>RNA support | **Rank 25**<br>RNA support | **Rank 21**<br>RNA support | 3/3 runs |
| GPRC5D | Not selected | **Rank 45**<br>RNA support not established | **Rank 95**<br>RNA support not established | 2/3 runs |
| SLAMF7 | Not selected | Not selected | Not selected | 0/3 runs |

Ranks refer to the candidate list. GPRC5D in Run 2 ranked 45 among candidates and 46 in the original full surface-gene list. RNA support refers to the external expression assessment, not a functional test.

#### B. Recovery at successive candidate-list cutoffs

| Candidates examined | Run 1 | Run 2 | Run 3 |
|---|---|---|---|
| Top 5 | 0/4 (0%) | 0/4 (0%) | 0/4 (0%) |
| Top 10 | 0/4 (0%) | 1/4 (25%) | 0/4 (0%) |
| Top 20 | 2/4 (50%) | 1/4 (25%) | 1/4 (25%) |
| Top 25 | 2/4 (50%) | 2/4 (50%) | 2/4 (50%) |
| Top 50 | 2/4 (50%) | 3/4 (75%) | 2/4 (50%) |
| Full candidate list | 2/4 (50%)<br>86 candidates | 3/4 (75%)<br>56 candidates | 3/4 (75%)<br>126 candidates |

Values are the number and percentage of the four reference antigens recovered. MM, multiple myeloma.
