## Supplementary Tables for "General Purpose AI for Public Data Analysis and Experimental Hypothesis Generation": Supplementary_Table_S6.pdf

### Supplementary Table S6. Additional lung relationships and tests in other donors

#### Blind analysis

Results in a second public lung dataset. Donor counts are shown as total tested, followed by IPF and control donors. Each question uses donors with enough cells of the relevant populations.

| ID | Question and cell populations | Donors Total (IPF / control) | Effect estimate (95% confidence interval) | q value | Outcome |
|---|---|---|---|---|---|
| B-L01 | Shared immunoglobulin RNA in AT2 and endothelial cells<br>AT2 / endothelial | 48<br>(22 / 26) | 0.723 (0.540 to 0.841)<br>Adjusted correlation, r | $3.22 \times 10^{-8}$ | Recurring background RNA |
| B-L02 | Shared heat-shock gene expression in monocytes and lymphoid cells<br>Monocyte / T and NK | 45<br>(20 / 25) | 0.927 (0.866 to 0.961)<br>Adjusted correlation, r | $2.66 \times 10^{-17}$ | Recurring stress RNA |
| B-L03 | Shared interferon-response gene expression in myeloid populations<br>Dendritic / macrophage | 0 tested<br>14 IPF eligible;<br>0 controls | Not tested<br>Adjusted correlation | Not applicable | Insufficient control cells |
| B-L04 | Shared antioxidant-related gene expression in AT2 and secretory cells<br>AT2 / secretory | 47<br>(25 / 22) | 0.830 (0.704 to 0.906)<br>Adjusted correlation, r | $2.67 \times 10^{-11}$ | Supported |
| B-L05 | Shared fatty-acid-oxidation gene expression in endothelial cells and macrophages<br>Endothelial / macrophage | 49<br>(22 / 27) | 0.283 (-0.015 to 0.535)<br>Adjusted correlation, r | 0.0376 | Supported, weaker effect* |
| B-L06 | Shared erythroid RNA in endothelial cells and macrophages<br>Endothelial / macrophage | 49<br>(22 / 27) | 0.590 (0.355 to 0.755)<br>Adjusted correlation, r | $1.51 \times 10^{-5}$ | Recurring background RNA |
| B-L07 | Rare epithelial cells coexpressing neuroendocrine markers<br>Epithelial | 70<br>(28 / 42) | 78 cells in 5 donors<br>Marker-positive cells | Not applicable | Cell identity supported; disease association not established |

A positive adjusted correlation means that donors with higher gene-set expression in one cell population also tended to have higher expression in the other, after accounting for diagnosis and technical variables. Recurring background or stress RNA may reflect sample processing or RNA originating outside the assigned cell population.

*The fatty-acid-oxidation result supported the predicted positive direction in a one-sided test, but its two-sided 95% confidence interval included zero and it did not remain significant with stricter cell filtering. The rare-cell search examined all 70 donors and detected 78 neuroendocrine-like cells in five donors.

q is the P value adjusted for multiple testing. Additional definitions are given on the next page.

### Supplementary Table S6. Additional lung relationships and tests in other donors

#### Context-informed analysis

Results in a second public lung dataset. Donor counts are shown as total tested, followed by IPF and control donors. Each question uses donors with enough cells of the relevant populations.

| ID | Question and cell populations | Donors Total (IPF / control) | Effect estimate (95% confidence interval) | q value | Outcome |
|---|---|---|---|---|---|
| C-L01 | Higher fibroblast CRYM RNA in IPF<br>Fibroblast | 31<br>(16 / 15) | 4.52 (2.10 to 9.94)<br>CRYM RNA rate ratio | $6.84 \times 10^{-4}$ | Supported |
| C-L02 | Higher endothelial glycolysis-related gene expression in IPF<br>Endothelial | 50<br>(22 / 28) | 0.445 (0.112 to 0.778)<br>Gene-set score difference | 0.0059 | Supported |
| C-L03 | Different IPF-associated damage-response RNA changes in AT2 cells and macrophages<br>AT2 / macrophage | 59<br>(25 / 34) | 1.553 (1.227 to 1.879)<br>Difference between cell populations | $1.21 \times 10^{-5}$ | Supported |
| C-L04 | Higher lymphoid MHC-II-related gene expression in IPF<br>Lymphoid | 51<br>(23 / 28) | 1.018 (0.644 to 1.392)<br>Gene-set score difference | $2.49 \times 10^{-6}$ | Supported |
| C-L05 | Shared antioxidant-related gene expression in epithelial and myeloid cells<br>Epithelial / myeloid | 63<br>(26 / 37) | 0.388 (0.144 to 0.587)<br>Adjusted correlation, r | 0.0020 | Supported |
| C-L06 | Shared oxidative-phosphorylation gene expression in myeloid and vascular cells<br>Myeloid / vascular | 51<br>(22 / 29) | 0.563 (0.326 to 0.733)<br>Adjusted correlation, r | $4.64 \times 10^{-5}$ | Supported |

Gene-set scores combine standardized expression of the genes specified in the deposited Part 3 external-validation records. Positive score differences indicate higher expression in IPF than controls. For C-L03, the estimate compares the disease-associated changes in AT2 cells and macrophages. A CRYM RNA rate ratio greater than 1 indicates higher fibroblast CRYM RNA abundance in IPF after accounting for total RNA measured.

q is the P value adjusted for multiple testing. Not applicable indicates that no q value was supplied for an untested question or a cell-identity count. Gene lists, detailed statistical results, and the additional comparisons and controls are provided in the deposited Part 3 external-validation records. AT2, alveolar type 2; IPF, idiopathic pulmonary fibrosis; NK, natural killer; MHC-II, major histocompatibility complex class II.

Computational repository: https://doi.org/10.5281/zenodo.23290610. The repository file index identifies the Part 3 external-validation records.
