## Supplementary Tables for "General Purpose AI for Public Data Analysis and Experimental Hypothesis Generation": Supplementary_Table_S10.pdf

### Supplementary Table S10. Related AI approaches for biological data analysis

#### Single-cell and spatial analysis

Comparison of 33 approaches, including the present study. Available workflow and testing details are included with each approach.

| Approach and source | Scope and workflow | Evaluation and testing |
|---|---|---|
| **CellAgent**<br>Specialized workflow/analysis agent<br>Conference publication; detailed evidence from supplied preprint<br>Supplied preprint; ICLR 2026 proceedings independently identified<br>Primary source | **Input:** User-provided scRNA-seq or spatial data and natural-language task<br>**Activity:** QC, clustering, annotation, batch correction, trajectory and spatial analysis<br>**Interface:** Conversational interface<br>**Setup:** Hosted page advertised; final-conference setup not fully verified<br>**Execution:** Planner, Executor and Evaluator; code execution through sc-Omni<br>**Data acquisition:** User-provided datasets in inspected examples | **Evaluation:** Multi-dataset benchmarks; iterative code debugging and result evaluation<br>**Human involvement:** Expert task scoring; 20-participant usability evaluation in supplied manuscript<br>**Experimental testing:** Not reported in inspected sections |
| **CellAtria**<br>Specialized workflow/analysis agent<br>Peer-reviewed article<br>npj Artificial Intelligence 2:8 (2026); published 16 January<br>doi:10.1038/s44387-025-00064-0 | **Input:** Article URL/PDF, repository identifiers or count matrices<br>**Activity:** Defined preprocessing, integration, annotation and report generation<br>**Interface:** Chatbot with logs, file browser and configuration controls<br>**Setup:** Docker deployment and model-provider configuration<br>**Execution:** Containerized LangGraph orchestration; CellExpress pipeline<br>**Data acquisition:** Document-derived GEO retrieval and preidentified repository downloads | **Evaluation:** Schema validation; configuration/provenance exports and HTML reports<br>**Human involvement:** Interactive demonstration plus autonomous dataset benchmark<br>**Experimental testing:** Not reported in inspected sections |
| **CellVoyager**<br>Specialized workflow/analysis agent<br>Peer-reviewed article<br>Nature Methods 23:749-759 (2026); online 17 March<br>doi:10.1038/s41592-026-03029-6 | **Input:** Single-cell object plus biological context and optionally prior analyses<br>**Activity:** Iterative hypothesis, code, execution, interpretation and replanning<br>**Interface:** Python/Jupyter workflow; generated notebook and report<br>**Setup:** Python environment, LLM access and prepared input data<br>**Execution:** Persistent Jupyter kernel with selected analysis packages<br>**Data acquisition:** Local single-cell object; optional literature background retrieval | **Evaluation:** CellBench analysis-proposal evaluation; preserved notebooks; explicit failed-code handling<br>**Human involvement:** Expert evaluation of case studies, including source-paper coauthors<br>**Experimental testing:** Not reported in inspected case-study sections |

### Supplementary Table S10. Related AI approaches for biological data analysis

#### Single-cell and spatial analysis (continued)

| Approach and source | Scope and workflow | Evaluation and testing |
|---|---|---|
| **scChat**<br>Conversational single-cell analysis<br>Peer-reviewed article<br>doi:10.1002/aic.70285 | **Input:** Single-cell data and successive biological questions<br>**Activity:** Plans and executes analyses with contextual follow-up | **Evaluation:** Annotation benchmarks and a glioblastoma immunotherapy reanalysis |
| **SpatialAgent**<br>Specialized spatial/workflow analysis agent<br>Preprint; peer review not established in audit<br>bioRxiv preprint v2; posted 12 August 2026<br>doi:10.1101/2025.04.03.646459 | **Input:** Spatial or single-cell data, reference images and natural-language tasks<br>**Activity:** Panel design, multimodal annotation, spatial analyses and hypotheses<br>**Interface:** Programmatic/notebook agent and scientist follow-up<br>**Setup:** Python environment and supported LLM API or local model<br>**Execution:** Plan-Act-Conclude; dynamic tools, reusable skills and saved artifacts<br>**Data acquisition:** Database/literature searches and provided datasets | **Evaluation:** Backend/skill ablations; numerical and visual checking; available traces<br>**Human involvement:** Autonomous benchmarks and illustrative scientist-guided analyses<br>**Experimental testing:** Prospective customized Xenium panel; researchers replaced 5 of top 100 selected genes |
| **OmniCellAgent**<br>Omics discovery agent<br>Preprint; not peer reviewed<br>doi:10.1101/2025.07.31.667797 | **Input:** Natural-language question and curated public omics resource<br>**Activity:** Selects datasets, analyzes expression and interprets evidence | **Evaluation:** Disease case studies and computational evaluations |
| **SCassist**<br>Workflow assistant / task-specific interpretation<br>Peer-reviewed Applications Note<br>Bioinformatics 41(8):btaf402 (2025); online 12 July<br>doi:10.1093/bioinformatics/btaf402 | **Input:** Seurat objects, computed metrics and enrichment results<br>**Activity:** Parameter recommendations, annotations, pathway interpretation and networks<br>**Interface:** R functions embedded in a Seurat workflow<br>**Setup:** R/package dependencies; commercial API key or local Ollama<br>**Execution:** User-managed R/Seurat plus selected LLM backend<br>**Data acquisition:** Prepared user Seurat data; enrichment/database context | **Evaluation:** Groundedness/semantic assessments; documented templates, package and tutorials<br>**Human involvement:** Eight senior/junior scientists rated reports from two reanalyses<br>**Experimental testing:** No new wet-lab validation reported in inspected evaluation |

### Supplementary Table S10. Related AI approaches for biological data analysis

#### Broader biomedical analysis and research platforms

| Approach and source | Scope and workflow | Evaluation and testing |
|---|---|---|
| **Biomni**<br>General biomedical agent<br>Peer-reviewed article; corrected supplementary material<br>Science 393:eadz4351 (2026); online 9 July; corrected supplement 21 August<br>doi:10.1126/science.adz4351 | **Input:** Natural-language biomedical tasks and diverse data inputs<br>**Activity:** Omics, literature, molecular design, protocol generation and automation<br>**Interface:** Python agent plus advertised no-code web interface<br>**Setup:** Hosted option; local environment and API keys for open implementation<br>**Execution:** Curated biomedical software/data environment with code execution<br>**Data acquisition:** Database APIs, downloaded data lake and user inputs | **Evaluation:** Task benchmarks, data/code archives; molecular predictions distinguished from experiments<br>**Human involvement:** Expert task reviews and cloning-protocol comparison<br>**Experimental testing:** B2M sgRNA cloning confirmed by Sanger; laboratory-automation demonstrations |
| **BioMedAgent**<br>General biomedical agent<br>Peer-reviewed article<br>Nature Biomedical Engineering (2026); published 30 March<br>doi:10.1038/s41551-026-01634-6 | **Input:** Natural-language questions and biomedical data<br>**Activity:** Omics, statistics, machine learning, visualization and pathology analysis<br>**Interface:** Natural-language task interface; code demos and official web platform advertised<br>**Setup:** Open implementation: Python, Redis, model API and supporting tools<br>**Execution:** Multi-agent planning/programming with tool learning and memory retrieval<br>**Data acquisition:** Curated benchmark inputs and tool-mediated data access | **Evaluation:** BioMed-AQA, external BixBench and accessible execution records<br>**Human involvement:** Benchmarks and expert/reference milestone evaluation<br>**Experimental testing:** Not reported in inspected study sections |
| **BioMaster**<br>Bioinformatics workflow agent<br>Peer-reviewed article<br>doi:10.1016/j.patter.2026.101611 | **Input:** Analysis task and data files<br>**Activity:** Plans, executes, checks and repairs analysis workflows | **Evaluation:** Multi-omics workflow benchmarks |
| **CARIBOU**<br>Configurable research-agent framework<br>Preprint; not peer reviewed<br>doi:10.64898/2026.05.25.727730 | **Input:** Analysis instructions, datasets and computational blueprints<br>**Activity:** Runs reproducible bioinformatics workflows in container environments | **Evaluation:** Task benchmarks, metadata reconstruction and single-cell case studies |

### Supplementary Table S10. Related AI approaches for biological data analysis

#### Broader biomedical analysis and research platforms (continued)

| Approach and source | Scope and workflow | Evaluation and testing |
| --- | --- | --- |
| **PromptBio**<br>Biomedical analysis platform<br>Preprint; not peer reviewed<br>doi:10.64898/2026.09.02.748774 | **Input:** Research task and biomedical data<br>**Activity:** Orchestrates planning and analysis with scientist oversight | **Evaluation:** Preprint platform description and case studies |
| **BioChatter**<br>Biomedical language-model framework<br>Peer-reviewed article<br>doi:10.1038/s41587-024-02534-3 | **Input:** Biological questions and connected biomedical resources<br>**Activity:** Supports natural-language access to tools and knowledge sources | **Evaluation:** Platform report |
| **The Virtual Biotech**<br>Therapeutic evidence synthesis<br>Peer-reviewed article<br>doi:10.1126/science.aeg6779 | **Input:** Genetics, cell-resolved data, clinical information and research questions<br>**Activity:** Coordinates agents to investigate therapeutic targets and development decisions | **Evaluation:** Published computational evaluations and target case studies |

### Supplementary Table S10. Related AI approaches for biological data analysis

#### Discovery and experimental feedback

| Approach and source | Scope and workflow | Evaluation and testing |
| --- | --- | --- |
| **Robin**<br>General biomedical discovery agent<br>Peer-reviewed article<br>Nature 655:497-505 (2026); online 19 May<br>doi:10.1038/s41586-026-10652-y | **Input:** Disease question followed by scientist-generated experimental data<br>**Activity:** Assay selection, candidate nomination, data analysis and iterative hypotheses<br>**Interface:** Multi-agent literature and data-analysis workflow<br>**Setup:** Framework/tool access; scientists perform experiments<br>**Execution:** Literature agents and experimental-data-analysis agent<br>**Data acquisition:** Literature retrieval and new assay data supplied by researchers | **Evaluation:** Iterative assay testing with archived trajectories/code<br>**Human involvement:** Scientists conducted laboratory experiments and returned data<br>**Experimental testing:** Candidate testing in retinal-cell phagocytosis models |
| **Co-Scientist**<br>Hypothesis generation<br>Peer-reviewed article<br>doi:10.1038/s41586-026-10644-y | **Input:** Scientist-defined research goal and literature<br>**Activity:** Generates, critiques and prioritizes research hypotheses | **Evaluation:** Scientist-led AML, liver-fibrosis and bacterial-evolution experiments |
| **BioPathfinder**<br>Evidence-guided hypothesis discovery<br>Preprint; not peer reviewed<br>doi:10.64898/2026.07.15.738646 | **Input:** Public studies, curated omics and a CAR-T objective<br>**Activity:** Links hypothesis generation with selected computational and laboratory tests | **Evaluation:** CAR-T repeated-stimulation and mouse xenograft experiments |
| **Kosmos**<br>Autonomous data-driven discovery<br>Preprint; not peer reviewed<br>doi:10.48550/arXiv.2511.02824 | **Input:** A supplied dataset and broad research objective<br>**Activity:** Alternates data analysis, literature search and hypothesis refinement | **Evaluation:** Independent analyses; cases with source findings withheld or unavailable |

### Supplementary Table S10. Related AI approaches for biological data analysis

#### Discovery and experimental feedback (continued)

| Approach and source | Scope and workflow | Evaluation and testing |
|---|---|---|
| **AI Sleep Co-Scientist**<br>Expert-guided physiological discovery<br>Preprint; not peer reviewed<br>doi:10.48550/arXiv.2607.25175 | **Input:** Large clinical and epidemiological sleep cohorts<br>**Activity:** Specialist agents develop hypotheses, preprocess signals and analyze data with expert review | **Evaluation:** Five case studies, including cross-cohort analyses |
| **Co-Scientist real-world extension**<br>Scientific research with experimental feedback<br>Preprint; not peer reviewed<br>doi:10.48550/arXiv.2608.26701 | **Input:** Research goals, laboratory constraints and experimental observations<br>**Activity:** Connects agent analysis with materials experiments and biological phenotype prediction | **Evaluation:** Materials studies and comparison with held-out biological measurements |
| **LabOS**<br>AI-assisted laboratory research<br>Preprint; not peer reviewed<br>doi:10.48550/arXiv.2510.14861 | **Input:** Research questions, functional screens and live laboratory video<br>**Activity:** Combines computational target analysis with visual guidance during experiments | **Evaluation:** NK-cell resistance assays, cell-fusion tests and stem-cell workflow demonstrations |
| **The AI Scientist**<br>Autonomous machine-learning research<br>Peer-reviewed article<br>doi:10.1038/s41586-026-10265-5 | **Input:** A computational research topic and experiment environment<br>**Activity:** Conducts machine-learning research cycles | **Evaluation:** Machine-learning evaluations |

### Supplementary Table S10. Related AI approaches for biological data analysis

#### Interpretation, representation models and data resources

| Approach and source | Scope and workflow | Evaluation and testing |
|---|---|---|
| **GeneAgent**<br>Task-specific gene-set interpretation agent<br>Peer-reviewed article<br>Nature Methods 22:1677-1685 (2025); online 28 July<br>doi:10.1038/s41592-025-02748-6 | **Input:** User-provided gene set<br>**Activity:** Biological-process naming and explanatory gene-set narratives<br>**Interface:** Programmatic pipeline<br>**Setup:** Python environment, model access and database APIs<br>**Execution:** Generation, database self-verification, modification and summarization<br>**Data acquisition:** Domain-database queries from gene symbols | **Evaluation:** Gene-set benchmarks, source-masking strategy, database evidence reports<br>**Human involvement:** Human checks of verification claims and study-specific interpretation<br>**Experimental testing:** Analysis of experimental gene sets; new causal testing not established by inspected sections |
| **GPTCelltype**<br>Cell annotation<br>Peer-reviewed article<br>doi:10.1038/s41592-024-02235-4 | **Input:** Marker genes from clusters<br>**Activity:** GPT-4 proposes cell-type names | **Evaluation:** Annotation benchmark |
| **CellWhisperer**<br>Transcriptome-language model<br>Peer-reviewed article<br>doi:10.1038/s41587-025-02857-9 | **Input:** Expression profiles and free-text queries<br>**Activity:** Links expression to language and supports interactive cell exploration | **Evaluation:** Annotation tests and human-development meta-analysis with marker follow-up |
| **scGPT**<br>Representation/foundation model<br>Peer-reviewed article<br>Nature Methods 21:1470-1480 (2024); online 26 February<br>doi:10.1038/s41592-024-02201-0 | **Input:** Tokenized cell-by-gene expression and condition information<br>**Activity:** Cell representations, annotation, integration, perturbation prediction and gene-network analysis<br>**Interface:** Python model and task-specific workflows<br>**Setup:** Model/software setup; compute appropriate to inference/fine-tuning<br>**Execution:** Generative pretrained transformer and downstream adaptation<br>**Data acquisition:** Prepared training/evaluation datasets | **Evaluation:** Pretraining/fine-tuning descriptions and public code/models<br>**Human involvement:** Task-specific computational evaluation<br>**Experimental testing:** No prospective wet-lab validation established in inspected methods |

### Supplementary Table S10. Related AI approaches for biological data analysis

#### Interpretation, representation models and data resources (continued)

| Approach and source | Scope and workflow | Evaluation and testing |
|---|---|---|
| **Minerva**<br>Genome-language-model discovery framework<br>Preprint; peer review not established in audit<br>bioRxiv preprint; supplied version dated 24 September 2026<br>doi:10.64898/2026.09.22.753630 | **Input:** Prokaryotic genomic sequences<br>**Activity:** Coevolutionary mining of noncoding sequence elements<br>**Interface:** Computational sequence-model framework<br>**Setup:** Model and sequence-analysis workflow setup<br>**Execution:** Genome-language-model interaction inference plus experiments<br>**Data acquisition:** Genome/metagenome sequence collections | **Evaluation:** Known-system recovery and perturbation-supported biochemical readouts<br>**Human involvement:** Investigator-designed biochemical characterization<br>**Experimental testing:** UG27 expression, catalytic mutants, PAGE and strand-specific DNA sequencing |
| **scBaseCount**<br>Public-data curation<br>Peer-reviewed article<br>doi:10.1016/j.cell.2026.08.025 | **Input:** Public single-cell repositories<br>**Activity:** Finds, organizes and standardizes reusable single-cell data | **Evaluation:** Published repository description |

### Supplementary Table S10. Related AI approaches for biological data analysis

#### Evaluation frameworks and the present study

| Approach and source | Scope and workflow | Evaluation and testing |
|---|---|---|
| **M3A**<br>Evaluation framework with agent execution harness<br>Preprint; peer review not established in audit<br>bioRxiv preprint v1; posted 9 June 2026<br>doi:10.64898/2026.06.04.729919 | **Input:** Prepared multi-omic data and staged biological-reasoning tasks<br>**Activity:** Annotation, gene-program hypotheses, regulatory analyses and biological synthesis<br>**Interface:** Autonomous and human-AI copilot experimental interfaces<br>**Setup:** Controlled Docker environment and model/tool configuration<br>**Execution:** Common initial environment reset between runs; persistent within-run state<br>**Data acquisition:** Prepared public cohorts and literature tools with source-study restrictions | **Evaluation:** Logged actions, external-cohort testing and blinded expert output review<br>**Human involvement:** Explicit autonomous, human-in-the-loop and human-on-the-loop comparisons<br>**Experimental testing:** No new wet-lab data generated in the study |
| **PromptBio-Bench**<br>Benchmark/evaluation framework<br>Preprint; peer review not established in audit<br>bioRxiv preprint v2 (2026); distinct from 194-task v1<br>doi:10.64898/2026.05.05.723092 | **Input:** Task capsules containing prompts, input files, reference answers and guidelines<br>**Activity:** 244 bioinformatics/data-science tasks in supplied v2<br>**Interface:** Evaluation code and task collection<br>**Setup:** Agent environments and evaluator configuration<br>**Execution:** Containerized agent executions with bounded task time<br>**Data acquisition:** Expert-curated inputs rather than open-ended dataset discovery endpoint | **Evaluation:** Structured file comparison and task-level execution logs<br>**Human involvement:** Expert task curation; computational output evaluation<br>**Experimental testing:** Not a wet-lab study |
| **POPPER**<br>Hypothesis-testing framework<br>Peer-reviewed conference paper<br>Primary source | **Input:** A proposed hypothesis and accessible observations<br>**Activity:** Translates claims into successive statistical tests | **Evaluation:** Computational evaluations across multiple domains |

### Supplementary Table S10. Related AI approaches for biological data analysis

#### Evaluation frameworks and the present study (continued)

| Approach and source | Scope and workflow | Evaluation and testing |
|---|---|---|
| **Single-cell agent benchmark**<br>Evaluation study<br>Peer-reviewed article<br>doi:10.1186/<br>s13059-026-03998-z | **Input:** Single-cell analysis tasks across models and agent frameworks<br>**Activity:** Measures task execution and robustness | **Evaluation:** 50 tasks with repeated-run, prompt and dataset analyses |
| **Present study**<br>Empirical use of general-purpose conversational and agentic interfaces<br>Author-review manuscript draft<br>Computational records<br>doi:10.5281/<br>zenodo.23290610 | **Input:** Biological prompts; selected or autonomously acquired public molecular datasets<br>**Activity:** Interactive hPSC hypothesis generation and laboratory testing; three PDAC and three MM autonomous runs; source-blinded lung reanalysis and external testing; broader public-data PDAC integration | **Evaluation:** Nonsignificant Apelin experiments; candidate recurrence and established-antigen recovery; 13 targeted external lung questions; contribution of multiple evidence types to PDAC ranking |

The present-study DOI links to computational records. PromptBio and PromptBio-Bench are separate entries. Missing details indicate that they were not included in the source comparison, not that a feature or test was absent from the original publication.
